# Satellite cell glucocorticoid receptor restrains myogenic commitment without mediating the effects of therapeutic glucocorticoids on muscle repair

**DOI:** 10.64898/2026.11.04.764308

**Authors:** Emilia Calvano, Aurore Bilger, Joe G. Rizk, Alicia Boule, Régis Lutzing, Erwan Grandgirard, Tao Ye, Céline Keime, Qingshuang Cai, Daniel Metzger, Sirine Souali-Crespo, Delphine Duteil

**Author notes:** Authors to whom correspondence should be addressed: Sirine Souali-Crespo and Delphine Duteil.

## Abstract

Glucocorticoids (GCs) exert profound effects on skeletal muscle, but the contribution of satellite cell (SatC) glucocorticoid receptor (GR) signaling to muscle regeneration remains poorly understood. Here, we investigated the role of GR in SatC homeostasis and assessed whether SatC GR mediates the effects of therapeutic GCs on muscle repair. Conditional deletion of GR in SatC reduced the number of PAX7-positive cells during regeneration in young mice, but did not substantially impair muscle repair in either young or aged animals. Transcriptomic and chromatin profiling revealed that GR occupies genes involved in stem cell fate and developmental signaling, with extensive GR binding already established in quiescent SatC. GR-deficient quiescent SatC showed reduced expression of *Yap1*, *Runx1* and *Foxo1*, together with increased *Myog* expression. Similarly, GR silencing in human myoblasts induced early myogenic differentiation markers without affecting proliferation, supporting a conserved role for GR in restraining myogenic commitment. We next examined the effects of two therapeutic GCs on muscle regeneration. Chronic prednisone treatment altered the transcriptional landscape of regenerating muscle in an age-dependent manner, without substantially affecting histological repair. In contrast, dexamethasone administration during the early inflammatory phase delayed regeneration, with persistent necrosis and increased collagen deposition. These effects were comparable in control and SatC-specific GR-deficient mice, indicating that they do not require GR in SatC. Together, these findings identify GR as a regulator of the quiescent, undifferentiated state of SatC, while showing that the effects of therapeutic GCs on muscle repair are largely independent of SatC GR. They further highlight the importance of treatment regimen and physiological context in determining the effects of GCs on skeletal muscle regeneration.

## Introduction

Muscle regeneration is a highly coordinated process that relies on muscle stem cells residing within skeletal muscle, known as satellite cells (SatC) (Relaix & Zammit, 2012). SatC are located beneath the basal lamina of muscle fibers and are characterized in their quiescent state by the expression of the paired-box transcription factor PAX7 (Seale et al., 2000). Quiescence is actively maintained by a specific transcriptional program, the disruption of which can trigger SatC activation and compromise the maintenance of the stem cell pool (Machado et al., 2017). Following injury, SatC exit quiescence, proliferate as myogenic progenitors and differentiate under the control of myogenic regulatory factors (MRFs), including MYF5, MYOD, myogenin (MYOG) and MRF4, which orchestrate the progression of the myogenic program (Zammit, 2017). A subset of activated SatC returns to quiescence, thereby replenishing the stem cell pool through self-renewal and ensuring the capacity for future rounds of regeneration.

Regeneration involves not only SatC but also multiple cell populations within the muscle niche, including fibro-adipogenic progenitors (FAPs), endothelial cells, pericytes and immune cells. Neutrophils are among the first immune cells to infiltrate injured muscle and contribute to the early inflammatory response (Ruffell et al., 2009; Saclier et al., 2013). Macrophages subsequently participate in tissue debris clearance and support SatC activation. Changes in macrophage functional states then contribute to myogenic differentiation and tissue remodeling (Dort et al., 2019; Tidball, 2017). With aging, muscle regenerative capacity declines as SatC function becomes impaired and the muscle niche develops a state of chronic inflammation (Moiseeva et al., 2023). Thus, aged muscle presents a distinct inflammatory environment that may influence the cellular and molecular mechanisms underlying regeneration.

Muscle regeneration also takes place within an endocrine environment shaped by glucocorticoids (GCs), steroid hormones produced by the adrenal cortex under the control of the hypothalamic-pituitary-adrenal (HPA) axis. Their secretion follows a circadian rhythm and increases in response to stress (Spies et al., 2014). GCs regulate a broad range of physiological processes, including metabolism, circadian rhythms and immune responses. Their actions are mediated by the glucocorticoid receptor (GR, NR3C1), which, upon ligand binding, translocates to the nucleus and regulates gene expression through several mechanisms. These include direct binding to glucocorticoid response elements (GREs) in DNA, leading to transcriptional activation or repression, and interactions with other transcription factors, such as AP-1 and NF-κB, which can modulate their transcriptional activity (Beato et al., 1996; Meijsing et al., 2009). Synthetic GCs, including prednisone and dexamethasone, are widely prescribed to treat inflammatory diseases and certain myopathies. However, prolonged exposure can cause substantial adverse effects, including skeletal muscle wasting. The effects of GCs on skeletal muscle are therefore context-dependent and may differ according to the physiological or pathological setting, the compound used and the timing and duration of exposure. For example, dexamethasone administration has been reported to prolong and exacerbate the necrotic and inflammatory phases following muscle injury, thereby impairing regeneration (Otrocka-Domagała et al., 2019). Conversely, other studies suggest that GCs can promote myogenic differentiation in cultured myoblasts (Fang et al., 2020). These apparently contrasting effects highlight the need to understand how GCs act on distinct cellular populations during muscle repair.

Although the effects of GCs on mature skeletal muscle fibers have been extensively investigated, their role in SatC remains poorly understood. In particular, it is unclear how GR contributes to the regulation of SatC identity and myogenic progression, and whether its chromatin-binding profile changes as SatC transition from quiescence to activation. Furthermore, whether the effects of therapeutic GCs on muscle regeneration are mediated directly by GR in SatC or indirectly through other cell populations in the muscle niche remains to be established. This question is particularly relevant in aged muscle, where chronic inflammation and age-related changes in the regenerative niche may modify the response to GC treatment.

To address these questions, we combined a mouse model of conditional GR deletion in SatC (GR^(i)sat−/−^) with complementary studies in human myoblasts. We characterized GR chromatin occupancy and gene expression in SatC isolated from uninjured and injured muscle. We also investigated muscle regeneration following injury in young and aged mice exposed to distinct GC regimens, including chronic prednisone administration and short-term dexamethasone treatment during the early pro-inflammatory phase. Our findings indicate that GR contributes to the maintenance of quiescent SatC identity by restraining myogenic commitment. However, the effects of therapeutic GC administration on muscle regeneration are largely independent of GR in SatC, implicating other cell populations in mediating these effects.

## Results

### GR deletion in satellite cells does not substantially impair regeneration

Immunostaining showed that GR is expressed in PAX7-positive SatC in uninjured muscle and, following injury, in SatC, IBA1-positive macrophages and COL4-positive fibro-adipogenic progenitors (FAPs) (**Fig. S1A**). To investigate the role of endogenous glucocorticoid signaling through GR in muscle regeneration, we generated mice in which GR is selectively deleted in SatC following tamoxifen administration (GR^(i)sat−/−^ mice). Nine-week-old mice carrying floxed *Nr3c1* alleles, which encode GR, and the *Pax7*-CreER^T2^ transgene were treated with tamoxifen (TAM) for 5 days to induce *Nr3c1* ablation in SatC (**Fig. S1B-D**). TAM-treated littermates lacking the *Pax7*-CreER^T2^ transgene served as controls. To evaluate the functional consequences of GR ablation on skeletal muscle regenerative capacity, we subjected young (15-week-old) control and GR^(i)sat−/−^ mice to acute tibialis anterior (TA) muscle injury via cardiotoxin (CTX) injection one month after gene ablation, and determined regeneration features of muscle repair at multiple days post-injury (dpi). The regeneration time course of control TA muscles from 2 to 28 dpi is shown in **Fig. S1E**.

Histological analysis of hematoxylin and eosin (H&E)-stained sections at 7 dpi revealed similar regenerative profiles in both genotypes, including centrally nucleated myofibers, necrotic areas and immune cell infiltration (**Fig. 1A**). Consistent with these observations, myofiber cross-sectional area (CSA) distribution was similar between control and GR^(i)sat−/−^ mice (**Fig. 1B**), indicating that GR deletion in SatC does not markedly impair overall muscle regeneration. Similarly, immunostaining for the macrophage marker F4/80 revealed comparable macrophage infiltration, while Sirius Red staining showed similar levels of fibrosis between control and knock-out individuals (**Fig. S1F-G**).

**Figure 1:**
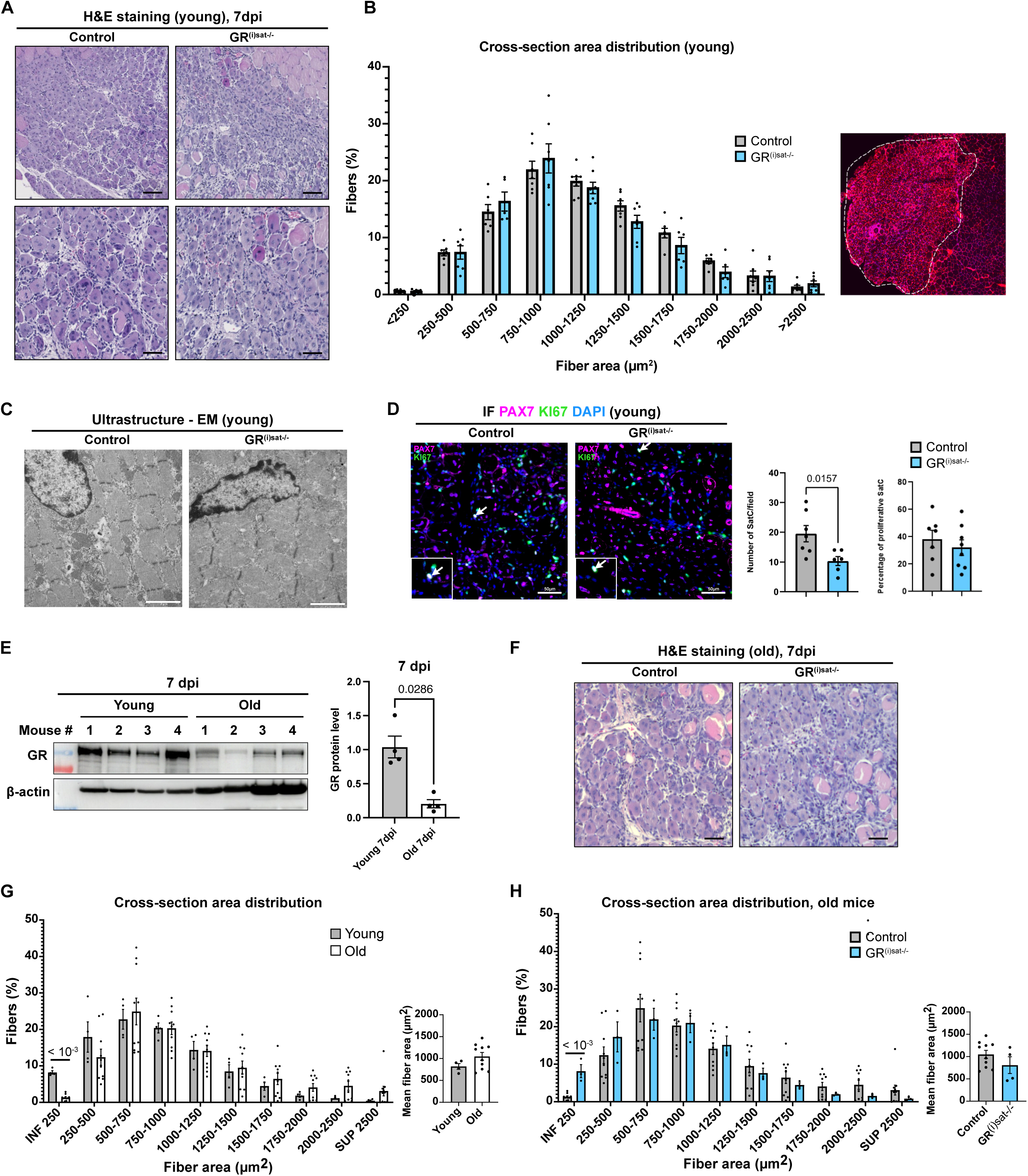
GR deletion in satellite cells does not impair regeneration. **(A)** Representative H&E staining of TA muscle sections from control and GR^(i)sat−/−^ mice at 7 dpi after CTX injury. Scale bars, 100 µm. **(B)** Myofiber cross-sectional area distribution in young control and GR^(i)sat−/−^ mice at 7 dpi; right, representative WGA (wheat germ agglutinin) staining of a TA section used for quantification, with the injured area outlined. Data are mean ± SEM. Statistical test used is Multiple unpaired t-test with Holm-Sidak post hoc correction. **(C)** Transmission electron microscopy of regenerating myofibers of young control and GR^(i)sat−/−^ mice at 7 dpi. **(D)** Immunofluorescent detection of PAX7 (magenta) and KI67 (green) at 7 dpi. Nuclei were counterstained with DAPI. White arrows indicate proliferative SatC. Quantification of satellite cell number (PAX7+) per field and of the proportion of proliferating (PAX7+KI67+) SatC. Scale bars, 50 µm. Data are mean ± SEM. Statistical test used was two-tailed Mann-Whitney test. **(E)** Western blot of GR in TA muscle of young and aged control mice at 7 dpi, with β-actin as loading control, and corresponding quantification. Data are mean ± SEM. Statistical test used was two-tailed Mann-Whitney test. **(F)** Representative H&E staining of aged control and GR^(i)sat−/−^ TA at 7 dpi. Scale bars, 50 µm. **(G)** Myofiber cross-sectional area distribution and mean fiber area in young and aged control mice at 7 dpi. Data are mean ± SEM. Statistical test used is Multiple unpaired t-test with Holm-Sidak post hoc correction. **(H)** Myofiber cross-sectional area distribution and mean fiber area in aged control and GR^(i)sat−/−^ mice at 7 dpi. Data are mean ± SEM. Statistical test used is Multiple unpaired t-test with Holm-Sidak post hoc correction.

Despite these broadly similar regenerative outcomes, ultrastructural analyses performed at 7 dpi revealed disorganized sarcomeres and disrupted nuclei in regenerating myofibers of GR^(i)sat−/−^ mice (**Fig. 1C**). In addition, the number of PAX7-positive SatC was reduced at 7 dpi, whereas the proportion of proliferating KI67-positive SatC was unchanged (**Fig. 1D**), suggesting that GR contributes to the maintenance of the SatC pool during regeneration.

To investigate whether SatC-specific GR deletion has more pronounced consequences in aged muscle, CTX injury was induced in 1-year-old control and GR^(i)sat−/−^ mice. As in the young cohort, tamoxifen-induced gene deletion had been initiated at 9 weeks of age (**Fig. S1B**). Of note, GR protein levels were markedly lower in whole regenerating muscles of aged control mice than in young ones (**Fig. 1E**). As expected, regenerating muscles of aged control mice displayed histological signs of impaired regeneration compared with young mice (**Fig. 1A and 1F**), consistent with a previous report (Rizk et al., 2026). Mean myofiber CSA, however, was similar between aged and young control mice, although young muscles contained a higher proportion of fibers smaller than 250 µm² (**Fig. 1G**). To determine whether GR loss in SatC contributes to this decline, we compared aged mutants with age-matched controls. At 7 dpi, H&E staining revealed no overt histological difference between aged GR^(i)sat−/−^ and control muscles (**Fig. 1F**), and mean myofiber CSA was similar between genotypes, although aged GR^(i)sat−/−^ muscles contained a higher proportion of fibers smaller than 250 µm² (**Fig. 1H**). Aged GR^(i)sat−/−^ muscles also showed no difference in macrophage infiltration or fibrotic deposition compared with age-matched controls (**Fig. S1H-I**), nor in interstitial space (**Fig. 1F**).

Overall, these results indicate that GR deletion in SatC reduces the SatC pool after injury in young mice but does not notably impair muscle regeneration in either young or aged mice.

### GR transcriptional activity in activated satellite cells

To elucidate the cell-specific GR-dependent molecular mechanisms governing SatC homeostasis during skeletal muscle regeneration, we performed SMART-seq on FACS-isolated SatC from TA muscles of young (15-week-old) control and GR^(i)sat−/−^ male mice at 7 dpi. Transcriptomic analysis revealed differential expression of 810 genes in GR-deficient SatC, including 323 up- and 487 down-regulated genes (**Fig. 2A and S2A**). *Nr3c1* transcripts were strongly reduced in GR-deficient SatC, confirming efficient gene deletion (**Fig. 2C**). Up-regulated genes were enriched in Gene Ontology (GO) terms related to stress response, cytoskeleton organization and developmental processes (**Fig. 2B**), and included *Hspb1*, *Tmod4* and *Itga5* (**Fig. 2C**). Conversely, GO enrichment analysis of the down-regulated gene set revealed a significant association with translation initiation, mRNA metabolism and the regulation of epithelial cell proliferation and differentiation (**Fig. 2B**), and this set included *Klf9*, *Notch2* and *Runx1* (**Fig. 2C**).

**Figure 2:**
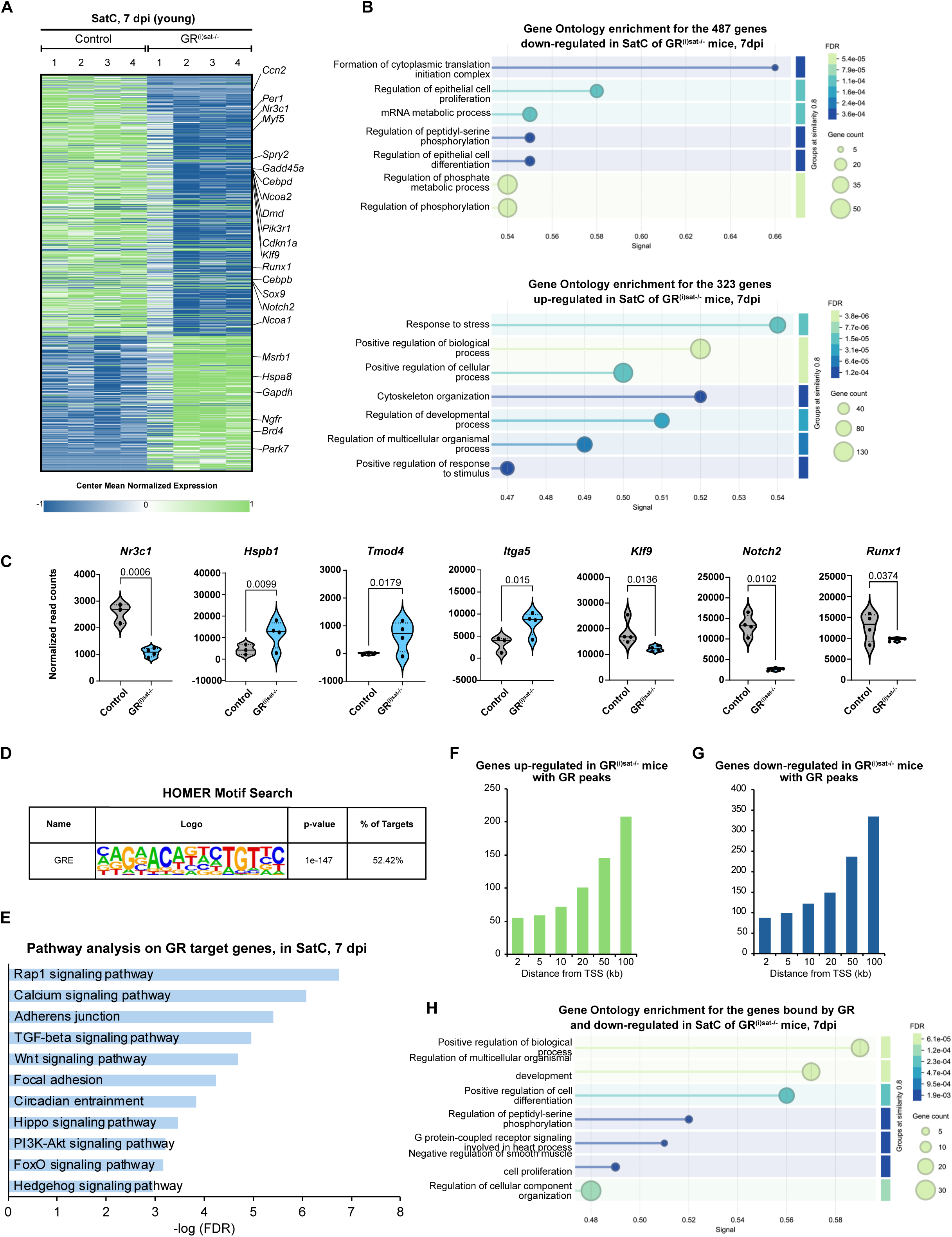
GR cistrome and transcriptome in activated satellite cells. **(A)** Heatmap showing the mean-centered normalized expression of differentially expressed genes from RNA-sequencing of FACS-sorted SatC isolated from TA muscles of control and GR^(i)sat−/−^ mice 7 dpi. **(B)** Gene Ontology enrichment for the 487 down-regulated (top) and 323 up-regulated (bottom) genes. **(C)** Violin plots showing normalized read counts of indicated genes from RNA-sequencing of FACS-sorted SatC isolated from TA muscles of control and GR^(i)sat−/−^ mice 7 dpi. P-values were obtained from DESeq2. **(D)** HOMER motif analysis of GR CUT&RUN peaks at 7 dpi. **(E)** KEGG pathway enrichment of GR-bound genes in SatC at 7 dpi. (**F, G**) Number of genes up- (F) and down-regulated (G) in GR^(i)sat−/−^ SatC carrying a GR peak within the indicated distance from the TSS. **(H)** Gene Ontology enrichment for genes both bound by GR and down-regulated in GR^(i)sat−/−^ SatC.

To identify genomic sites bound by GR, we isolated SatC from TA muscles of control mice by FACS at 7 dpi and performed CUT&RUN assays using a GR-specific antibody. Bioinformatic analysis identified 11,416 GR-binding peaks, associated with 6,582 genes. Peaks were predominantly located in intronic and intergenic regions (**Fig. S2B**), and motif analysis showed that 52.4% of them contained a canonical GRE (**Fig. 2D**). Integrative analysis with the previously characterized cistrome of H3K4me2, a hallmark of active promoters and enhancers (Rizk et al., 2026), revealed that GR is recruited to 5,323 transcribed genes (**Fig. S2C**). Pathway enrichment analysis of GR-bound genes uncovered a strong association with *Rap1* and calcium signaling, adherens junctions and TGF-β signaling, as well as with key developmental and regenerative pathways, including the Wnt (*Wnt3a*, *Wnt4*, *Fzd1*, *Fzd5*, *Ctnnb1*), Hippo (*Stk3*, *Lats2*, *Wwtr1*, *Ptpn14*, *Limd1*), FoxO (*Foxo1*, *Foxo3*, *Akt2*, *Akt3*, *Pik3r1*) and Hedgehog (*Shh*, *Ihh*, *Ptch1*, *Gas1*, *Gli3*) signaling cascades (**Fig. 2E**). Representative GR peaks at the *Fzd2* and *Foxo3* loci are shown in **Fig. S2E**. The overlap between our cistrome and transcriptome analyses revealed that GR was bound to 17% of the up-regulated (54/323) and 20% of the down-regulated (99/487) genes in SatC of GR^(i)sat−/−^ mice (**Fig. S2D**). For more than half of these genes, GR peaks lay within a 100-kb window around the transcription start site (TSS) (**Fig. 2F-G**). Genes that were both bound by GR and down-regulated in GR^(i)sat−/−^ SatC were enriched in developmental processes and positive regulation of cell differentiation (**Fig. 2H**), whereas no significant enrichment was detected for the up-regulated set.

Altogether, GR binds a limited fraction of the genes deregulated in its absence, mostly at distal sites at 7 dpi, consistent with limited phenotypic abnormalities, and genes that are both bound and down-regulated are enriched in developmental processes.

### Chronic prednisone treatment elicits distinct transcriptional responses in young and aged regenerating muscle

To investigate the impact of sustained GC exposure on muscle regeneration, we applied a chronic prednisone treatment, initiated 7 days before injury and maintained for 7 days after injury (1 mg/kg/day in young and 0.1 mg/kg/day in aged mice; **Fig. 3A**). Using this experimental procedure, we assessed regenerative outcomes following cardiotoxin-induced injury in young and aged control mice. In young mice, histological analysis revealed misshapen fibers in prednisone-treated muscles (**Fig. 3B**). However, quantification showed no significant difference in mean myofiber CSA, fiber size distribution, or macrophage infiltration between prednisone-treated and vehicle-treated muscles (**Fig. 3C and S3A**). Mean myofiber CSA was also similar between vehicle- and prednisone-treated mice at 28 dpi, with only a lower proportion of 2,000-2,500 µm² fibers in prednisone-treated muscles (**Fig. S3B**).

**Figure 3:**
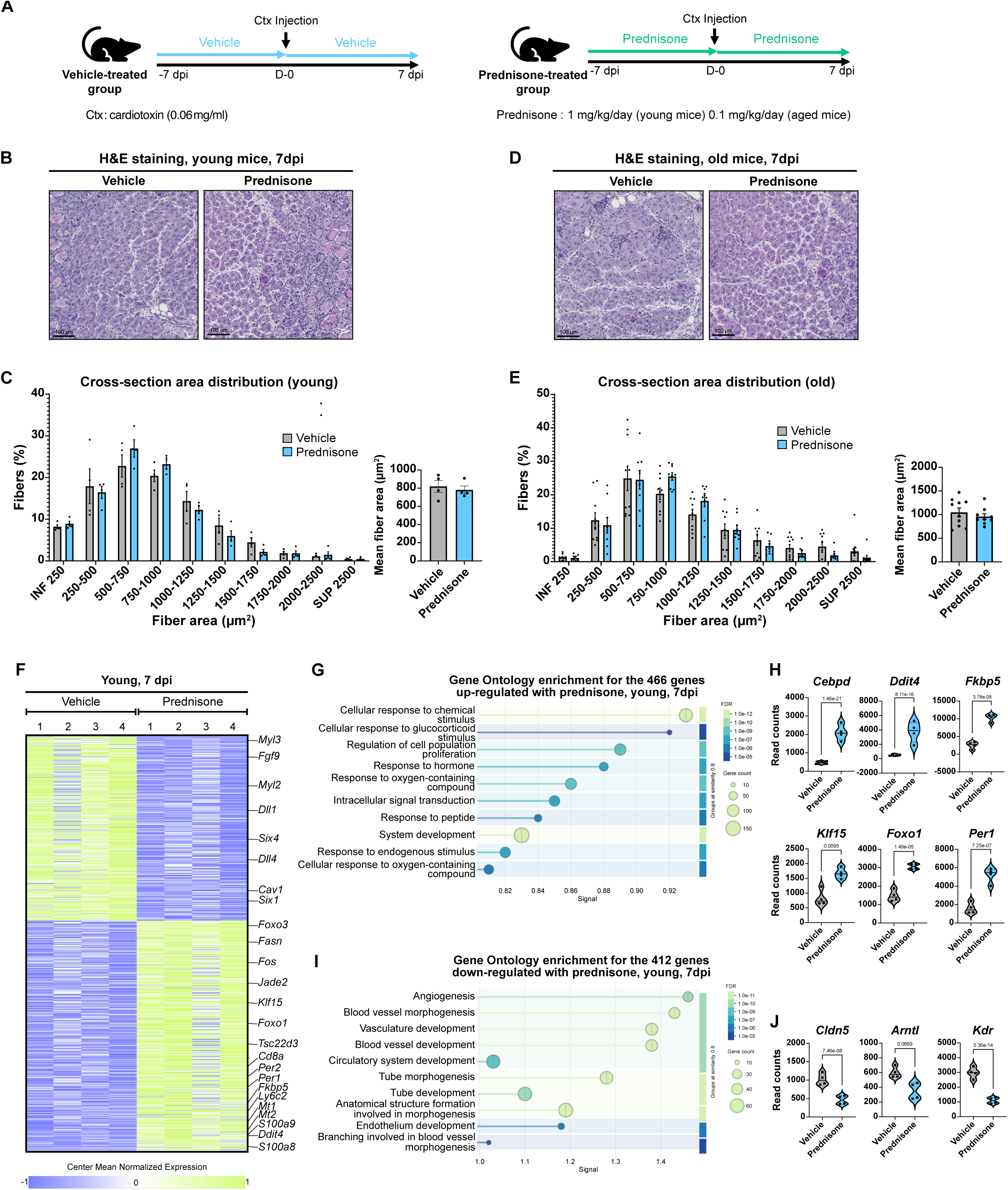
Age-dependent response to chronic prednisone. (**A**) Experimental design. Prednisone (1 mg/kg/day in young, 0.1 mg/kg/day in aged mice) or vehicle (water containing 1.5% EtOH) was administered from 7 days before CTX injury (0.06 mg/ml) of the TA until 7 dpi. (**B-E**) Representative H&E staining of TA sections from young (B) and aged (D) mice at 7 dpi, and corresponding myofiber cross-sectional area distribution and mean fiber area in young and aged mice (C, E) Scale bars, 100 µm. Data are mean ± SEM. Statistical test used is Multiple unpaired t-test with Holm-Sidak post hoc correction. (**F**) Heatmap of the 878 genes differentially expressed in injured TA of young mice treated with prednisone or vehicle. (**G-J**) Gene Ontology enrichment for the 466 up-regulated (G) and 412 down-regulated (I) genes, and violin plots showing normalized read counts of representative up- (H) and down-regulated (J) genes. P-values were obtained from DESeq2.

In aged mice, prednisone treatment affected neither mean fiber size nor cross-sectional area distribution (**Fig. 3D-E**) and did not modify macrophage infiltration (**Fig. S3A**) or interstitial fibrosis (**Fig. S3C**). In contrast, lipid droplet accumulation was significantly increased in prednisone-treated muscles (**Fig. S3D**), suggesting an altered fate of FAPs, which can give rise to adipocytes during regeneration.

To gain insight into the molecular basis underlying these divergent responses, we performed RNA sequencing on injured tibialis anterior muscles from young and aged mice treated with vehicle or prednisone. Transcriptomic analyses revealed 412 down- and 466 up-regulated genes differentially expressed in young mice after prednisone treatment (**Fig. 3F and S3E**). Up-regulated genes were enriched in cellular response to GC and hormone stimuli, including canonical GR targets (*Fkbp5*, *Ddit4*, *Tsc22d3*, *Klf15*, *Cebpd*, *Foxo1*) and circadian regulators (*Per1*, *Per2*) (**Fig. 3F-H**). Conversely, down-regulated genes were enriched in angiogenesis (*Cldn5*, *Cav1*, *Dll4*, *Kdr*) and developmental processes (*Six1*, *Six4*, *Dll1*, *Fgf9*) (**Fig. 3F, I-J**). Expression of the core clock gene *Arntl* also tended to decrease (**Fig. 3J**).

Prednisone led to the increased expression of 284 genes in aged mice, while repressing 287 transcripts (**Fig. S3F-G**). In contrast with young mice, the 287 down-regulated genes identified in aged mice after prednisone treatment were associated with inflammation and antigen presentation (*Cd68*, *Cd74*, *Cd40* and the MHC class II transactivator *Ciita*) and extracellular matrix organization (*Fn1*, *Acta2*) (**Fig. S3G-I**). Conversely, the 284 up-regulated genes belonged to cellular respiration, oxidative phosphorylation, electron transport chain and mitochondrion organization (*Ppargc1a*, *Sucla2*, *Cox6c*, *Ugp2*) (**Fig. S3G**, **S3J-K**). Notably, no canonical GC signature was detected in aged muscles, as none of the direct GR target genes induced in young mice (*Ddit4*, *Tsc22d3*, *Klf15*) were found among the genes up-regulated upon prednisone treatment (**Fig. S3L**).

Thus, the transcriptional response to chronic prednisone differed markedly between young and aged mice. In young mice, prednisone induced a canonical GR signature and repressed angiogenic and developmental genes, whereas in aged mice it induced no canonical GR signature, repressed inflammatory and extracellular matrix genes, and induced genes involved in mitochondrial respiration.

### Satellite cell GR deletion barely modifies regeneration under chronic prednisone

To determine whether GR in SatC impacts muscle regeneration under chronic prednisone, we next analyzed regenerative outcomes in GR^(i)sat−/−^ mice under the same chronic treatment conditions.

Following quantification, mean fiber area was not affected by prednisone in either young or aged mice (**Fig. S4A-B**). In young mice, the fiber size distribution was comparable across the four conditions, except for a higher proportion of fibers in the 500-750 µm² class in prednisone-treated GR^(i)sat−/−^ mice (**Fig. 4A**). Furthermore, macrophage infiltration (F4/80) did not differ between genotypes or treatments (**Fig. 4C**).

**Figure 4:**
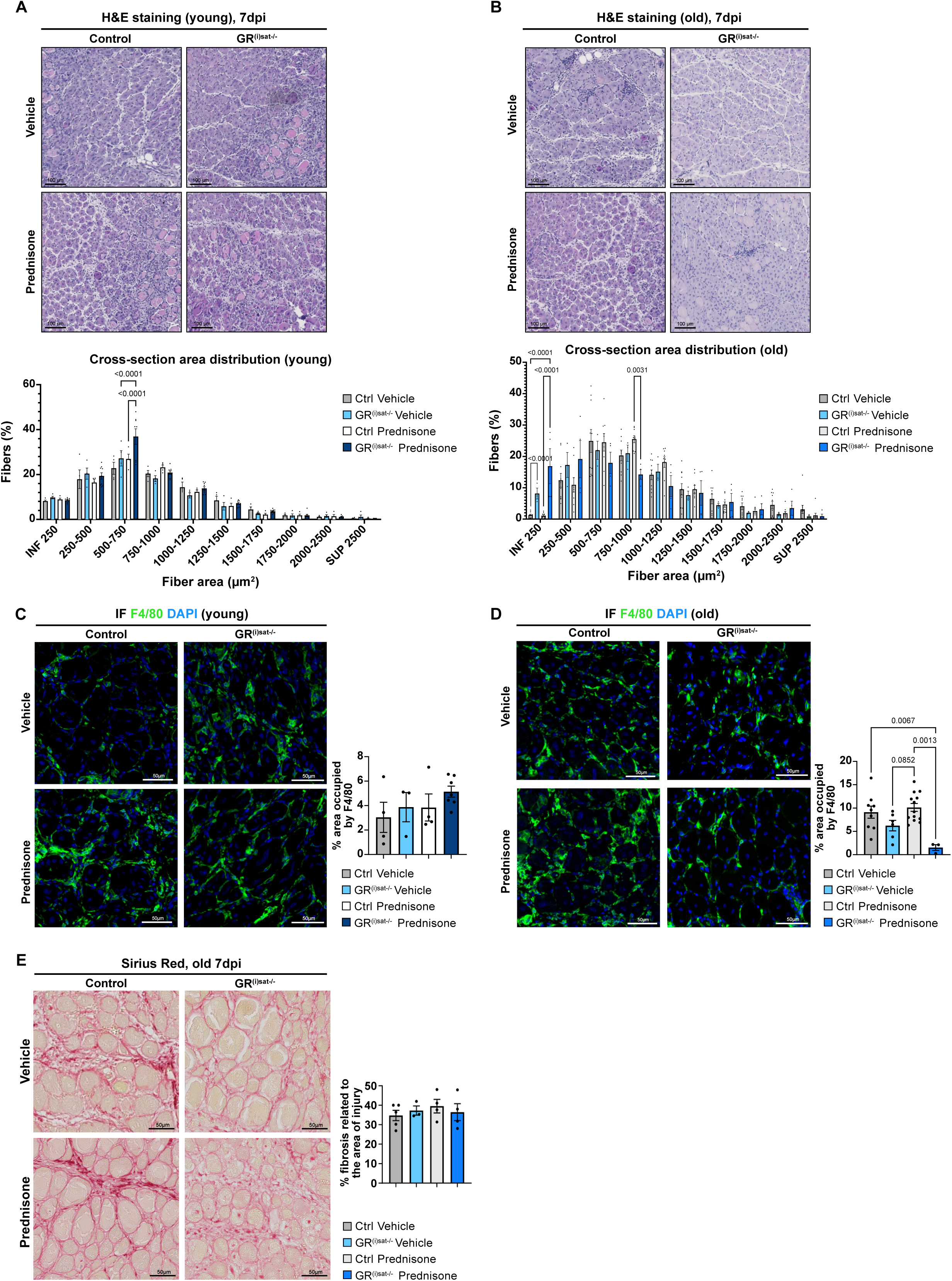
Prednisone effects are largely independent of satellite cell GR. (**A, B**) Representative H&E staining of TA sections from young (A) and aged (B) control and GR^(i)sat−/−^ mice treated with vehicle or prednisone at 7 dpi, and corresponding myofiber cross-sectional area distribution. Scale bars, 100 µm. Data are mean ± SEM. Statistical test used is Two-way ANOVA with Tukey post-hoc correction. (**C, D**) Immunofluorescent detection of F4/80 and corresponding quantification in young (C) and aged (**D**) mice. Nuclei were counterstained with DAPI. Scale bars, 50 µm. Data are mean ± SEM. Statistical test used is Kruskal-Wallis. (**E**) Sirius Red staining and quantification of fibrosis in aged mice. Scale bars, 50 µm. Data are mean ± SEM. Statistical test used is Kruskal-Wallis. Prednisone was administered at 1 mg/kg/day (young mice) or 0.1 mg/kg/day (aged mice).

In aged mice, H&E staining revealed no marked differences in the untreated groups, but GR^(i)sat−/−^ muscles treated with prednisone exhibited a reduced inflammatory infiltrate compared with controls, with no difference in mean fiber size, although these mice showed a higher proportion of small fibers (**Fig. 4B and S4B**). Consistent with this observation, the area occupied by F4/80+ macrophages was significantly reduced in GR^(i)sat−/−^ muscles treated with prednisone, compared with vehicle- and prednisone-treated controls (**Fig. 4D**). Interstitial fibrosis (Sirius Red) did not differ across the four conditions (**Fig. 4E**). Thus, except for a reduced macrophage infiltration in prednisone-treated aged mutants, SatC GR deletion did not modify regenerative outcomes under chronic prednisone in young and aged mice. This macrophage phenotype suggests that SatC GR may influence macrophage recruitment in aged muscle, through mechanisms that remain to be identified.

### Early dexamethasone delays regeneration independently of GR in satellite cells

To further assess the impact of GC exposure during the early phases of muscle regeneration, we treated young control and GR^(i)sat−/−^ mice with dexamethasone (1 mg/kg/day) during the initial pro-inflammatory phase (day 0 to day 2 post-injury; **Fig. S5A**) and analyzed regenerative outcomes at 5, 7, 14 and 28 dpi. Under these conditions, dexamethasone-treated mice exhibited markedly impaired muscle regeneration, as evidenced by a high number of persistent necrotic fibers and by the appearance of emerging myofibers, reminiscent of the early damage phase, compared with vehicle-treated cohorts at 5 dpi (**Fig. S5B**), consistent with the delayed regeneration previously reported upon early GC exposure (Souali-Crespo et al., 2026). At 14 and 28 dpi, no overt histological difference was observed between vehicle- and dexamethasone-treated muscles in either genotype (**Fig. S5B**).

By 7 dpi, histological analyses revealed that whereas inflammation had largely resolved in vehicle-treated mice, TA muscles of dexamethasone-treated control and GR^(i)sat−/−^ mice displayed sustained necrosis and an exacerbated stromal response (**Fig. 5A**), indicating that transient GC exposure during the early inflammatory phase delays the regenerative process. Of note, dexamethasone shifted the fiber size distribution toward smaller fibers in both genotypes, and mean myofiber CSA decreased upon treatment, reaching significance in GR^(i)sat−/−^ muscles but not in controls, with no difference between genotypes after treatment (**Fig. 5B**), indicating that the dexamethasone-induced reduction in fiber size does not require SatC GR. Macrophage infiltration (F4/80) was higher in dexamethasone-treated GR^(i)sat−/−^ muscles than in vehicle-treated controls, although the difference did not reach significance when compared with either dexamethasone-treated controls or vehicle-treated GR^(i)sat−/−^ mice (**Fig. 5C**). Collagen deposition was also increased upon dexamethasone treatment in both genotypes (**Fig. 5D**). These results show that dexamethasone given during the early inflammatory phase delays regeneration, and that this effect does not require GR in SatC.

**Figure 5:**
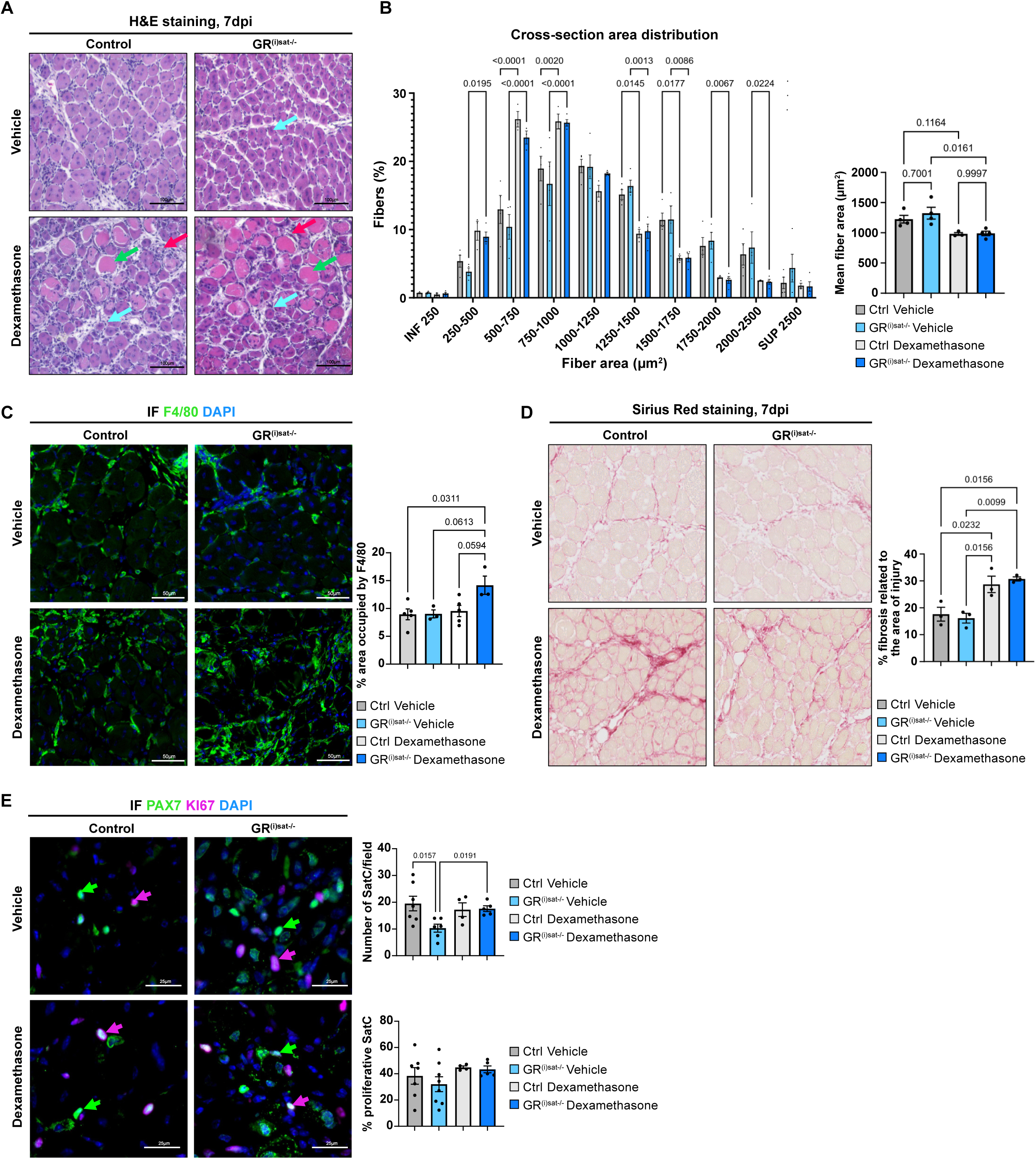
Dexamethasone impairs regeneration independently of satellite cell GR. **(A)** Representative H&E staining of TA sections at 7 dpi from control and GR^(i)sat−/−^ mice treated with vehicle or dexamethasone from day 0 to day 2 post-injury. Cyan arrows indicate newly formed fibers, green arrows necrotic fibers, red arrows inflammatory infiltrate. Scale bars, 100 µm. **(B)** Myofiber cross-sectional area distribution and mean cross-sectional area. Data are mean ± SEM. Statistical test used is Two-way ANOVA with Tukey post-hoc correction (CSA distribution) or Kruskal-Wallis (mean CSA). **(C)** Immunofluorescent detection of F4/80 and corresponding quantification of the area occupied by macrophages. Nuclei were counterstained with DAPI. Scale bars, 50 µm. Data are mean ± SEM. Statistical test used is Kruskal-Wallis. **(D)** Sirius Red staining and corresponding quantification of fibrosis over the injured area. Scale bars, 50 µm. Data are mean ± SEM. Statistical test used is Kruskal-Wallis. **(E)** Immunofluorescent detection of PAX7 (green) and KI67 (magenta), and corresponding quantification of satellite cell number per field and of the proportion of proliferating SatC. Green arrows indicate PAX7+ SatC, magenta arrows PAX7+ KI67+ proliferative SatC. Nuclei were counterstained with DAPI. Scale bars, 25 µm. Data are mean ± SEM. Statistical test used is Kruskal-Wallis. Dexamethasone was administered at 1 mg/kg/day.

Consistent with the reduced SatC pool previously observed in regenerating GR^(i)sat−/−^ muscles at 7 dpi (**Fig. 1D**), SatC numbers were significantly decreased in vehicle-treated GR^(i)sat−/−^ muscles. This difference was no longer detected upon dexamethasone treatment, GR^(i)sat−/−^ SatC numbers being comparable to controls (**Fig. 5E**). Importantly, the lower SatC number in vehicle-treated GR^(i)sat−/−^ muscles was not associated with alterations in SatC proliferation during regeneration, as comparable proportions of KI67-positive SatC were observed across all experimental conditions (**Fig. 5E**), indicating that GR signaling in SatC is dispensable for their proliferative response during acute inflammatory activation. These observations therefore point to a potential function of GR in controlling SatC states in muscle, prompting us to investigate its role in maintaining the quiescent SatC compartment.

### GR occupies stemness-associated loci in quiescent satellite cells

To explore the cellular context in which GR signaling may control SatC state, we took advantage of our previously published single-cell RNA sequencing at day 5 post-injury, a time point encompassing multiple SatC states along the regenerative trajectory (Rizk et al., 2026). In agreement with our immunofluorescence analysis, *Nr3c1* (encoding GR) was expressed in the various muscle cell populations (**Fig. 6A-C**). Unsupervised clustering identified four distinct SatC populations corresponding to quiescent, activated, dividing and differentiated states (**Fig. 6D and S6A-B**). Notably, *Nr3c1* was markedly enriched in the quiescent SatC cluster, whereas its expression was lower in activated, dividing and differentiated populations (**Fig. 6E-F**). The enrichment of GR expression within the quiescent SatC compartment suggests a potential role for GC signaling in the regulation of SatC homeostasis under basal conditions. To assess the consequences of GR loss in this context, we next profiled the transcriptome of SatC isolated from uninjured control and GR^(i)sat−/−^ muscles, together with the GR cistrome of SatC isolated from uninjured control muscles.

**Figure 6:**
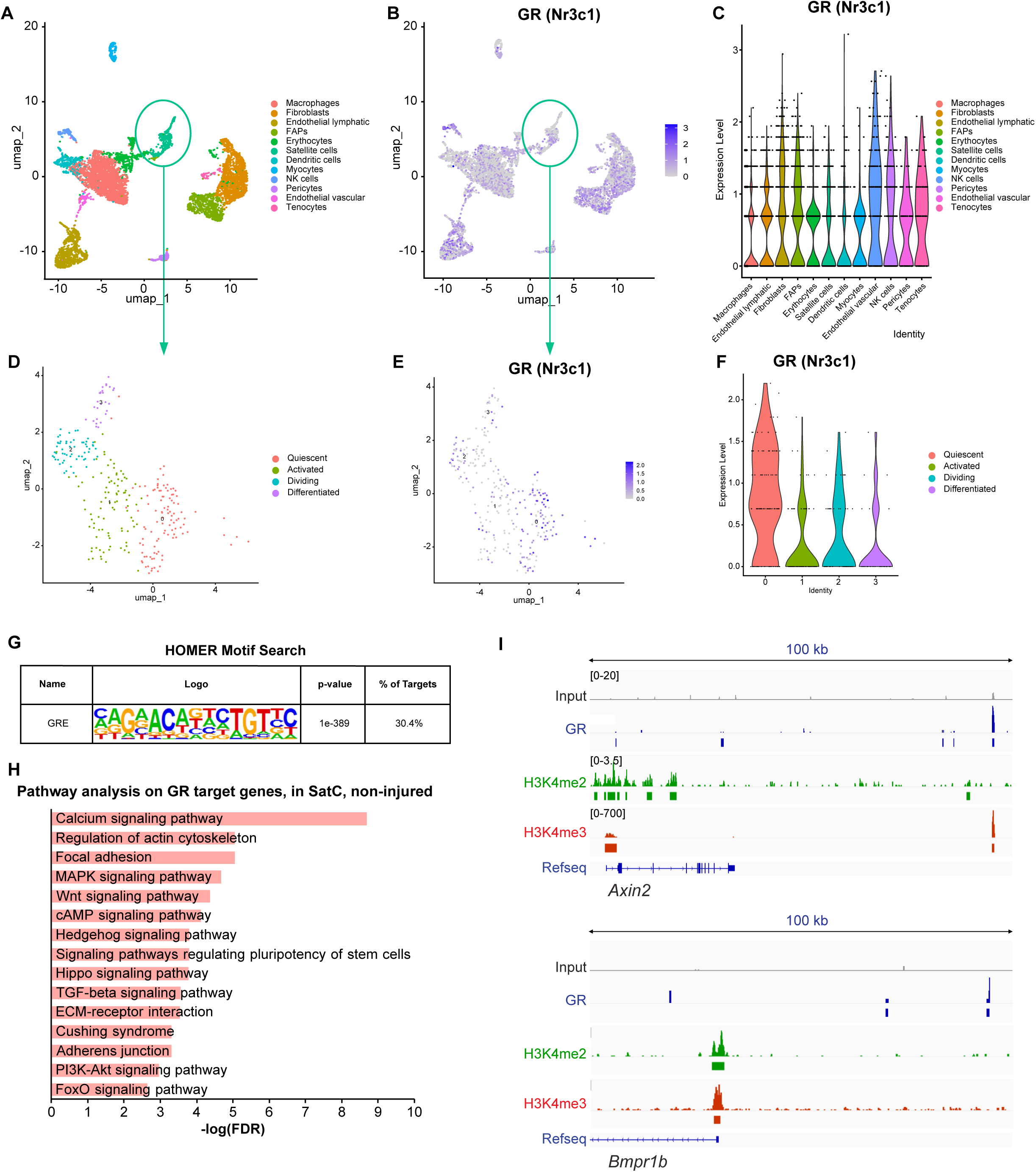
GR marks quiescent satellite cells. **(A)** Uniform manifold approximation and projection (UMAP) visualization of single-cell RNA-seq from regenerating muscle at 5 dpi (Rizk et al., 2026), colored by cell population. **(B)** Visualization of *Nr3c1* expression projected on the same UMAP. **(C)** Violin plot of *Nr3c1* transcript levels in the indicated cell populations identified from 5 dpi TA of wild-type mice. **(D)** UMAP visualization of SatC reclusters from 5 dpi TA, colored by cluster identity. SatC population were sub-clustered into quiescent, activated, dividing and differentiated states. **(E)** Visualization of *Nr3c1* expression projected on the satellite cell sub-clusters. **(F)** Violin plot of *Nr3c1* expression per SatC sub-cluster (0 = quiescent, 1 = activated, 2 = dividing, 3 = differentiated). **(G)** HOMER motif analysis of GR CUT&RUN peaks in SatC from uninjured muscle. **(H)** KEGG pathway enrichment of GR-bound genes in uninjured SatC. **(I)** GR, H3K4me2 and H3K4me3 localization at the *Axin2* and *Bmpr1b* loci in uninjured SatC.

CUT&RUN analysis identified 41,829 GR binding peaks, primarily localized to intronic and intergenic regions (**Fig. S6C**), associated with 13,808 genes. Motif analysis using HOMER revealed that 30.4% of GR peaks contained a canonical GRE (**Fig. 6G**), a lower proportion than in SatC isolated at 7 dpi (52.4%, **Fig. 2D**). Among the 17,617 genes marked by H3K4me2 in quiescent SatC (Rizk et al., 2026), 10,084 were bound by GR. Functional enrichment analysis of GR-bound genes uncovered a significant enrichment in pathways controlling stem cell fate, including Wnt, Hedgehog, Hippo, TGF-β and FoxO signaling and signaling pathways regulating pluripotency of stem cells, as well as pathways governing cellular responses to extracellular cues such as calcium, MAPK and PI3K-Akt signaling (**Fig. 6H**). Representative GR binding sites in uninjured SatC are shown at the *Axin2* (Wnt pathway) and *Bmpr1b* (TGF-β/BMP pathway) loci (**Fig. 6I**), and at the Hippo pathway components *Limd1* and *Dlg1* (**Fig. S6F**). To further assess the cell state-dependent role of GR, we compared CUT&RUN profiles of SatC isolated from uninjured muscle and from regenerating muscle at 7 dpi. Most GR-bound genes identified at 7 dpi were also bound in uninjured SatC (4,848 of 6,582 genes, 74%), whereas a large fraction of the genes bound in quiescent SatC were not detected at 7 dpi (8,960 of 13,808 genes, 65%; **Fig. S6D-E**). Thus, most genes bound by GR in activated SatC are already bound in quiescent SatC.

In uninjured muscle, SMART-seq profiling of freshly isolated SatC identified 1,600 transcripts differentially expressed between GR^(i)sat−/−^ and control mice (727 down- and 873 up-regulated; **Fig. 7A-B**). Intersecting this signature with the GR cistrome established in uninjured SatC (13,808 genes) showed that 701 of these transcripts (44%, 389 of the up- and 312 of the down-regulated genes) carry at least one GR binding site (**Fig. 7A**). GO enrichment analysis of this subset revealed a strong association with biological processes involved in morphogenesis, cell migration and cellular organization, including anatomical structure morphogenesis, regulation of cell migration and motility, cell projection organization and regulation of cell shape. In addition, enrichment was observed for signaling pathways controlling cell fate and developmental programs, such as transmembrane receptor protein tyrosine kinase signaling, system development and multicellular organism development (**Fig. 7C**). Consistently, individual examination of this dataset showed that the stemness-associated regulators *Yap1*, *Runx1* and *Foxo1* were down-regulated in GR^(i)sat−/−^ SatC, whereas the differentiation marker *Myog* was up-regulated and expression of the GR target gene *Ddit4* was unchanged (**Fig. 7D**). These data suggest a role for GR in maintaining the quiescent, undifferentiated state of the satellite cell pool.

**Figure 7:**
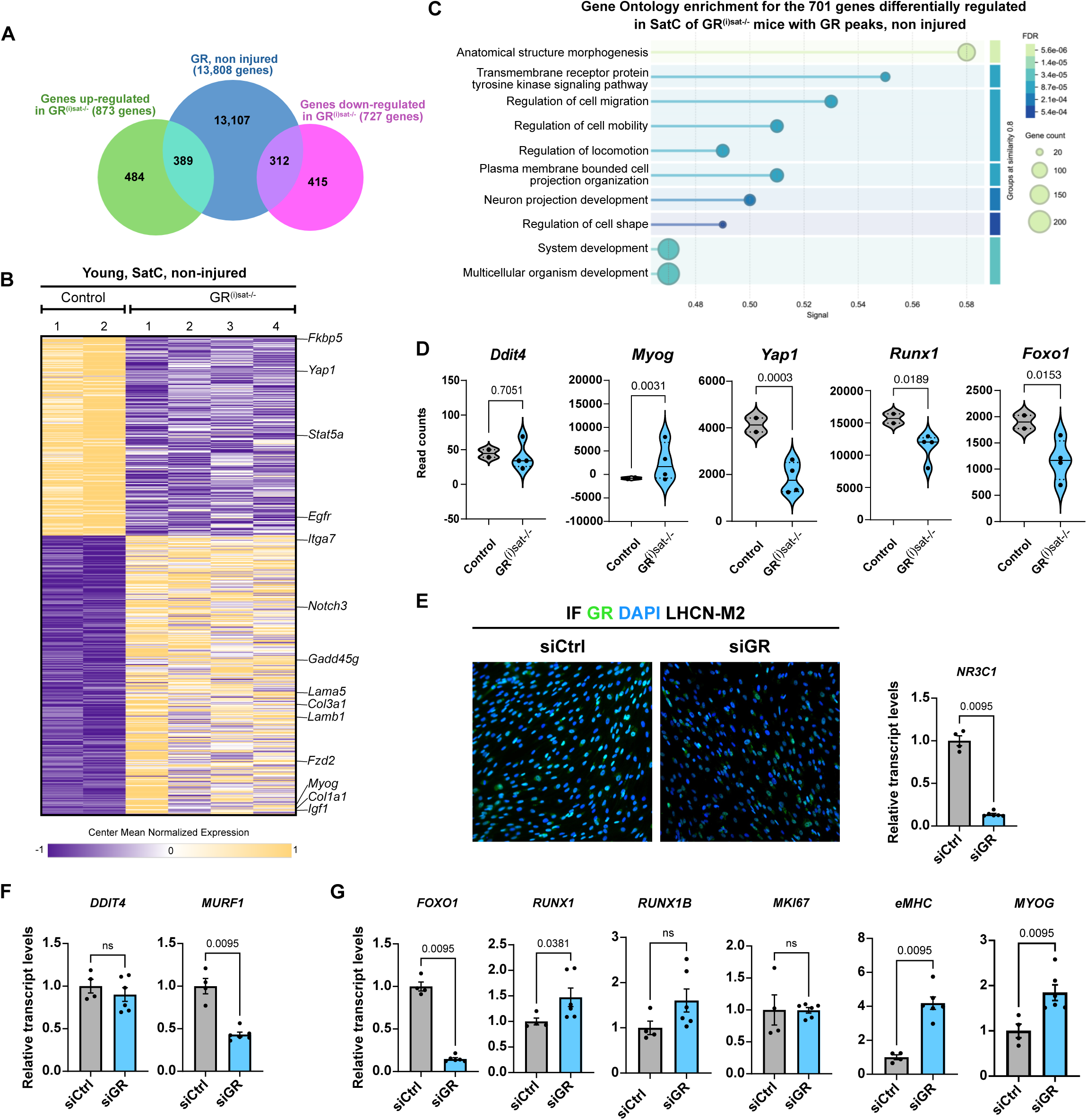
GR loss promotes early differentiation markers in uninjured satellite cells and human myoblasts. **(A)** Overlap between GR-bound genes in uninjured SatC (13,808 genes) and transcripts up- (873) or down-regulated (727) in GR^(i)sat−/−^ SatC from uninjured muscle. **(B)** Heatmap of the 1,600 gene differentially expressed in SatC from uninjured muscle of control and GR^(i)sat−/−^ mice. **(C)** Gene Ontology enrichment for the 701 differentially expressed genes with GR peak. P-values were obtained from DESeq2. **(D)** Violin plots showing normalized read counts of *Ddit4*, *Myog*, *Yap1*, *Runx1* and *Foxo1*. **(E)** Immunofluorescent detection of GR (green) of LHCN-M2 myoblasts transfected with control (siCtrl) or GR-targeting (siGR) siRNA, and *NR3C1* mRNA levels. **(F)** mRNA levels of the GR target genes *DDIT4* and *TRIM63* (MURF1). **(G)** mRNA levels of *FOXO1*, *RUNX1*, *RUNX1B*, *MKI67*, *MYH3* (eMHC) and *MYOG*. mRNA levels were normalized to *RPLP0*. Data are mean ± SEM. Statistical test used was two-tailed Mann-Whitney test.

### GR depletion promotes early myogenic differentiation markers in human myoblasts

To investigate whether the role of GR in maintaining the undifferentiated state of muscle progenitors is conserved in humans, we used LHCN-M2 myoblasts, an immortalized cell line derived from human pectoralis major muscle (Zhu et al., 2007), expressing *NR3C1* transcripts and GR protein (**Fig. 7E**).

To assess the role of GR in human muscle progenitors, we silenced its expression using a GR-targeting siRNA, with a scrambled siRNA as control, in charcoal-treated growth medium lacking dexamethasone, and analyzed the cells 72 h after transfection under non-differentiating conditions. GR transcript levels were reduced by approximately 85%, and the loss of GR protein was confirmed by immunostaining (Fig. 7E). Expression of the GR target gene *TRIM63* (MURF1) was reduced by half, whereas *DDIT4* expression was not significantly modified (**Fig. 7F**). Together, these results confirm the efficiency of GR silencing.

We next examined markers of the progenitor state. Among stemness-associated genes, *FOXO1* expression was almost abolished upon GR knockdown, whereas *RUNX1* was increased; its isoform *RUNX1B* showed a comparable trend that did not reach significance (Fig. 7G). The proliferation marker *MKI67* was unaffected (Fig. 7G). In contrast, the differentiation markers *MYH3* (eMHC) and *MYOG* were markedly induced (Fig. 7G). Taken together, in undifferentiated human myoblasts, GR knockdown increased the expression of early differentiation markers, as observed in murine SatC, without changing *MKI67* expression. In contrast, *RUNX1* was regulated in opposite directions in human myoblasts and murine SatC.

Collectively, these findings support a conserved role for GR in restraining the expression of early myogenic differentiation markers in murine SatC and human myoblasts, although the regulation of individual stemness-associated genes is not fully conserved between species.

## Discussion

Glucocorticoids are widely prescribed drugs that exert profound effects on skeletal muscle, yet the contribution of SatC GR signaling to muscle regeneration remains poorly understood. Here, we show that SatC GR helps maintain the quiescent, undifferentiated state of muscle stem cells but is largely dispensable for muscle repair. GR occupies stemness-associated loci in quiescent SatC, and its deletion is associated with reduced expression of *Yap1*, *Runx1* and *Foxo1*, together with increased *Myog* expression. Consistent with these findings, GR silencing in human myoblasts increases the expression of early myogenic differentiation markers without affecting proliferation. Furthermore, the effects of therapeutic GCs on muscle regeneration depend on the treatment regimen and are largely independent of SatC GR, as chronic prednisone modifies the muscle transcriptome in an age-dependent manner, whereas early dexamethasone treatment delays regeneration in both control and mutant mice.

Despite the reduced number of PAX7-positive SatC observed in GR-deficient muscles at 7 days post-injury (dpi), overall regeneration was largely preserved in both young and aged mice. The proportion of proliferating SatC was unchanged, indicating that the reduced SatC pool cannot be attributed to a detectable proliferation defect during regeneration. Whether this reduction reflects altered differentiation, survival or return to quiescence remains unresolved. Our findings are consistent with those of Rajgara et al. (2023), who also reported a reduced SatC pool and premature exit from quiescence following GR deletion, despite preserved muscle regeneration. Together, these studies support a role for GR in maintaining SatC identity, rather than a strict requirement for GR in regenerative myogenesis.

This function parallels the role of the androgen receptor (AR) in SatC described by Rizk et al. (2026), highlighting the contribution of nuclear hormone receptors to the regulation of muscle stem cell state. Although both receptors may influence SatC homeostasis, their downstream transcriptional programs and physiological functions should not be assumed to be identical. In our study, GR deletion altered the expression of stemness-associated regulators and increased *Myog* expression in murine SatC. Similarly, GR silencing in human myoblasts increased MYOG and MYH3 expression without affecting MKI67. However, the regulation of individual stemness-associated genes, notably RUNX1, differed between species, suggesting that the broader role of GR in restraining early myogenic commitment may be conserved despite differences in specific transcriptional responses.

Our cistromic analyses further suggest that GR participates in establishing the transcriptional landscape of SatC before injury. GR expression was enriched in quiescent SatC, and most genes bound by GR at 7 dpi were also associated with GR binding in uninjured muscle. In quiescent SatC, GR occupied loci associated with Wnt, Hippo, FoxO, Hedgehog and TGF-β signaling, pathways involved in stem cell maintenance and fate determination. These observations suggest that GR may help maintain the quiescent state through the regulation of multiple developmental and signaling networks. Although the number and distribution of identified binding sites differed between quiescent and activated SatC, these data do not establish a global loss of GR occupancy following activation.

The effects of exogenous GCs on muscle repair differed substantially from the consequences of SatC-specific GR deletion. Chronic prednisone treatment altered the transcriptome of regenerating muscle without substantially changing its histological appearance. In young mice, prednisone induced a canonical GC-responsive transcriptional signature and repressed genes associated with angiogenesis and development. In aged mice, it instead reduced the expression of inflammatory and extracellular matrix-associated genes and increased the expression of genes involved in mitochondrial respiration. These age-associated differences must be interpreted in light of the tenfold lower prednisone dose administered to aged mice for tolerability reasons. Nevertheless, the transcriptional findings indicate that the response to chronic GC exposure differs between the two experimental cohorts. Importantly, SatC GR deletion had little impact on the regenerative response to prednisone, apart from reduced macrophage infiltration in treated aged mutants. The significance and mechanism of this isolated phenotype remain to be established.

By contrast, dexamethasone administered during the first three days after injury markedly delayed regeneration, with persistent necrosis, increased collagen deposition and a shift toward smaller myofibers at 7 dpi. These effects occurred in both control and GR-deficient mice, indicating that SatC GR is not required for the deleterious effects of early dexamethasone exposure. This finding is consistent with the study by Souali-Crespo et al. (2026), which also supports a role for GC actions outside SatC in the regulation of muscle repair. Together, these observations suggest that the effects of dexamethasone are not primarily mediated by GR in SatC or, as reported in the myeloid compartment, by GR in myeloid cells. Rather, GC treatment may influence several cellular components of the regenerative niche.

The different outcomes of prednisone and dexamethasone likely reflect several factors, including treatment duration, timing relative to injury, compound-specific pharmacological properties and potentially differential engagement of glucocorticoid and mineralocorticoid receptors. Our experiments were not designed to isolate the contribution of each factor. Nevertheless, the pronounced delay observed after early dexamethasone exposure, compared with the limited histological effects of chronic prednisone, emphasizes that GC effects on muscle repair depend on the treatment context. Importantly, the delayed regeneration observed at 7 dpi should not be interpreted as permanent impairment, as muscle morphology was reported to recover at later time points in the study by Souali-Crespo et al. (2026), consistent with our observations at 14 and 28 dpi (**Fig. S5B**).

In conclusion, our findings identify SatC GR as a regulator of the quiescent, undifferentiated state of muscle stem cells, while demonstrating that it is largely dispensable for muscle regeneration under the conditions tested. Conversely, the effects of therapeutic GCs on muscle repair are strongly dependent on the treatment regimen and are mediated predominantly through mechanisms that do not require SatC GR. Understanding how GCs act on the different cellular populations within the muscle niche may help distinguish their therapeutic and detrimental effects on skeletal muscle.

## Materials & Methods

### Mouse housing

Mice were housed under a 12-h light/dark cycle, with standard rodent chow (2800 kcal/kg, Usine d’Alimentation Rationnelle, Villemoisson-sur-Orge, France) and water *ad libitum*. Breeding and maintenance of mice were performed according to institutional guidelines in an accredited animal facility, in compliance with French and EU regulations on the use of laboratory animals for research. All procedures were approved by the local ethics committee (Com’Eth, Strasbourg, France) and authorized by the French Research Ministry (MESR) according to Directive 2010/63/EU (authorization numbers 48090-2024022308468680 and 23477-2019121617036403).

### Generation of GR^(i)sat−/−^ mice

We generated a mouse line in which the glucocorticoid receptor (GR) is selectively inactivated in muscle satellite cells (SatC). Mice carrying floxed *Nr3c1* alleles (GR^L2/L2^, in which exon 3 is flanked by two loxP sites) (Tronche et al., 1999) and the *Pax7-CreER*^T2^ transgene (Murphy et al., 2011) were maintained on a C57BL/6J background. To delete *Nr3c1* in SatC, 9-week-old *Pax7-CreER^T2^*/GR^L2/L2^ males and their GR^L2/L2^ littermates received intraperitoneal injections of tamoxifen (1 mg/mouse/day) for five consecutive days. The resulting mutant mice are referred to as GR^(i)sat−/−^ mice, and tamoxifen-treated GR^L2/L2^ littermates were used as controls. Primers used for genotyping are listed in Table S1.

### Mouse treatments

For the cardiotoxin-induced injury model, young (15-week-old) and aged (1-year-old) male mice were anesthetized with 2.5% isoflurane using a SomnoSuite device, and 20 µl of 25 µM cardiotoxin (CTX; Latoxan, Valence, France, L8102) in phosphate-buffered saline (PBS) were injected bilaterally into the tibialis anterior (TA) muscles.

Prednisone (vitalis) was dissolved at 20 mg/ml in ethanol (EtOH) and diluted in water to 0.3 mg/ml for young mice and to 0.03 mg/ml for aged mice. It was administered by oral gavage (100 µl) at 1 mg/kg in young mice and 0.1 mg/kg in aged mice. Treatment started 7 days before CTX injection and was given daily until 7 days post-injury (dpi). Control mice received the vehicle (water containing 1.5% EtOH) by oral gavage on the same schedule.

Dexamethasone (Dex, Sigma, D1756) was dissolved at 20 mg/ml in EtOH, diluted to 0.3 mg/ml in sunflower oil and injected intraperitoneally at 1 mg/kg immediately after CTX injection (0 dpi) and again at 1 and 2 dpi. Control mice received intraperitoneal injections of the vehicle (sunflower oil containing 1.5% EtOH) on the same schedule.

Mice were euthanized by cervical dislocation at the indicated time points. TA muscles were immediately collected, weighed, and either frozen in liquid nitrogen for biochemical analyses or processed for histology and immunofluorescence.

### Histology

TA muscles were fixed overnight in 4% PFA at 4 °C, washed in 1× PBS, dehydrated and embedded in paraffin. Transverse sections of 5 µm were cut through the middle of the lesion and collected on slides. For hematoxylin and eosin staining, rehydrated sections were stained in filtered hematoxylin (Harris, 26041-05) for 3 min and 1% eosin Y (Sigma, 17372-87-1) for 3 min, then dehydrated and mounted. Collagen was revealed by Sirius Red staining on paraffin sections, performed by the histology platform of the Institut de Génétique et de Biologie Moléculaire et Cellulaire (IGBMC).

### Immunofluorescence on muscle sections

Deparaffinized TA sections underwent antigen retrieval in citrate unmasking solution (Cell Signaling, 14746) in a pressure cooker for 20 min, were cooled, and washed in PBS-0.1% Tween (PBST). Sections were left overnight at 4 °C in a humidified chamber with primary antibodies in PBST. Mouse or rabbit IgGs were used as controls. The next day, after three PBST washes, sections were incubated 1 h at room temperature in the dark with secondary antibody (Table S2). Slides were mounted in DAPI-Fluoromount-G (Invitrogen, 00-4959-52) and imaged on a Leica DM4000B epifluorescence microscope with a Hamamatsu ORCA-Flash camera, keeping exposure times constant across conditions.

All quantifications were performed in Fiji (ImageJ). Sarcolemmal boundaries of individual fibers were delineated with wheat germ agglutinin (WGA; Invitrogen, W32464; Table S2). Cross-sectional areas (CSAs) were measured with a custom Fiji macro developed by the IGBMC Imaging Center, which segments fibers based on the intensity and continuity of the WGA signal. The lesion area was delineated manually on the basis of DAPI and WGA staining. For F4/80, an intensity threshold was applied to retain specific signal and exclude background, and the F4/80^+^ area was normalized to the lesion area. Sirius Red-positive area was quantified with the QuPath pixel classifier and expressed relative to the injured area. PAX7^+^ SatC and KI67^+^PAX7^+^ cells were counted manually.

### Transmission electron microscopy

TA muscles collected at 7 dpi were processed as described (Rizk et al., 2026). Briefly, samples (1 mm^2^) were fixed overnight at 4 °C in 2.5% glutaraldehyde and 2.5% PFA in 0.1 M cacodylate buffer (pH 7.4), post-fixed in 1% osmium tetroxide for 1 h at 4 °C, dehydrated through graded ethanol and propylene oxide, and embedded in Epon 812. Ultrathin sections (70 nm) were contrasted with uranyl acetate and lead citrate and imaged at 70 kV on a Morgagni 268D electron microscope equipped with a MegaView III camera (Soft Imaging System).

### FACS isolation of satellite cells

SatC were isolated from 15-week-old male control and GR^(i)sat−/−^ mice, either uninjured (all the muscles of the limbs) or at 7 dpi (both TA). The muscle stromal vascular fraction was prepared as previously described (Ghaibour et al., 2023). Briefly, hindlimb muscles were dissected, minced and digested for 1 h at 37 °C under agitation in 1.25 U/ml Dispase II (Stemcell Technologies, 07913) and 500 µg/ml collagenase type I (Thermo Fisher Scientific, 17018029). The cell suspension was filtered through 70 µm cell strainers (Corning Life Sciences) to remove debris. Mononucleated cells were pelleted (400 g, 5 min) and resuspended in FACS buffer (phenol red-free DMEM with 4.5 g/l glucose, 2% bovine serum albumin and 2 mM EDTA) (Rizk et al., 2026). Cells were stained for 1 h on ice with a fixable viability dye (BD, 565388; 1:400) and with the following antibodies, all at 1:100: CD45-PE (eBioscience, 12-0451-83), CD11b-PE-Cy7 (eBioscience, 25-0112-82), CD31-PE (eBioscience, 12-0311-82), TER119-PE (BD, 553673) and ITGA7-FITC (MBL, K0046-4). After exclusion of debris and doublets on forward scatter (FSC) and side scatter (SSC), hematopoietic cells (CD45^+^, CD11b^+^ or TER119^+^) and endothelial cells (CD31^+^) were gated out, and SatC were sorted as the ITGA7^+^ fraction on a BD FACSAria. For RNA extraction, SatC were sorted into the lysis buffer of the RNeasy Plus Micro Kit (Qiagen, 74034). For CUT&RUN, sorted cells were processed directly. Data were analyzed with FlowJo as described (Ghaibour et al., 2023).

### CUT&RUN and cistrome analysis

CUT&RUN (Skene et al., 2018) was performed on FACS-isolated SatC as described (Ghaibour et al., 2023). Cells were pelleted (500 g, 10 min, 4 °C) and lysed for 20 min on ice in nuclear extraction buffer (20 mM HEPES-KOH pH 7.9, 10 mM KCl, 0.5 mM spermidine, 0.1% Triton X-100, 20% glycerol, protease inhibitors). Isolated nuclei were washed and bound to concanavalin A-coated beads (BioMag Plus Concanavalin A, Polysciences, 86057-3) for 10 min at 4 °C. After additional washes, nuclei-bead complexes were incubated overnight at 4 °C under gentle agitation with antibodies against GR (Proteintech, 24050-1-AP), H3K4me2 (Active Motif, 39141), H3K27ac (Active Motif, 39133) or H3K4me3 (Abcam, ab1012), all at 1:100 and each in a separate reaction. Normal rabbit IgG (Santa Cruz, sc-2027) was used as a negative control. Nuclei-bead complexes were then collected and incubated with protein A-micrococcal nuclease (pA-MNase; IGBMC; final concentration 700 ng/ml) for 1 h at 4 °C under constant agitation. Digestion was initiated by adding 3 µl of 100 mM CaCl_2_ (final concentration 2 mM in 150 µl; Skene et al., 2018) and carried out for 30 min on ice. The reaction was stopped with stop buffer (200 mM NaCl, 20 mM EDTA, 4 mM EGTA, 50 µg/ml RNase A, 40 µg/ml glycogen, 25 pg/ml yeast spike-in DNA), followed by 20 min at 37 °C. DNA was purified, and libraries were generated as described (Ghaibour et al., 2023).

Libraries were sequenced on an Illumina NextSeq 2000 as paired-end 50 bp reads and mapped to the mm10 reference genome with Bowtie2 v2.3.4.3 (Langmead & Salzberg, 2012). Only uniquely mapped, deduplicated reads were retained. Reads overlapping ENCODE blacklist regions (v2; Amemiya et al., 2019) were removed with BEDTools (Quinlan & Hall, 2010), and the remaining fragments were split by size: fragments shorter than 120 bp (sub-nucleosomal, used for transcription factors) and fragments longer than 150 bp (nucleosomal, used for histone marks).

Bigwig files were generated with bamCoverage (deepTools 3.3.0; Ramírez et al., 2016; --normalizeUsing RPKM --binSize 20), and raw bedGraph files with genomeCoverageBed (BEDTools v2.26.0). Peaks were called with SEACR (Meers et al., 2019) and MACS2, using IgG samples as controls (−keepall), and peaks with an FDR < 0.05 were retained.

Genome-wide signal profiles were visualized in IGV (http://software.broadinstitute.org/software/igv/; Thorvaldsdóttir et al., 2013) or Figeno (Sollier et al., 2024). Peaks were annotated and motifs searched with HOMER v4.11 (Heinz et al., 2010; findMotifsGenome.pl, mm10, GC-matched background). In de novo motifs, R denotes a purine (G or A) and Y a pyrimidine (T or C). Peaks were assigned to genomic features (promoter/TSS, 5Z UTR, exon, intron, 3? UTR, TTS and intergenic regions) with HOMER, based on RefSeq annotation and the distance to the nearest TSS.

Venn diagrams were generated with Venny 2.1.0 (https://bioinfogp.cnb.csic.es/tools/venny/) or InteractiVenn (Heberle et al., 2015). Pathway analysis was performed with WebGestalt using over-representation analysis (ORA), with a false discovery rate (FDR) below 0.05 as significance threshold. Clustering analyses were performed with seqMINER (Ye et al., 2011) and deepTools (computeMatrix --skipZeros; plotHeatmap). Genomic intersections and sequence extractions were carried out with BEDTools: getfasta was used to extract FASTA sequences from BED files, and intersectBed to identify overlapping genomic regions.

### RNA extraction and SMART-seq analysis on FACS-isolated SatC

RNA was isolated from FACS-isolated SatC of 15-week-old male control (GR^L2/L2^) and GR^(i)sat−/−^ mice, uninjured or at 7 dpi, with the RNeasy Plus Micro Kit (Qiagen, 74034) according to the manufacturer’s instructions, and RNA integrity was assessed on a Bioanalyzer. SMART-seq was performed on n = 2 control and n = 4 GR^(i)sat−/−^ for uninjured, and on n = 4 control and n = 4 GR^(i)sat−/−^ biological replicates for 7 dpi, each corresponding to SatC sorted from an individual mouse.

Library preparation was performed by the GenomEast platform (IGBMC). Full-length cDNA was generated from 250 pg to 2 ng of total RNA with the SMART-Seq mRNA Kit (Takara Bio Europe, Saint-Germain-en-Laye, France) according to the manufacturer’s instructions, with 12 PCR cycles of cDNA amplification by SeqAmp polymerase. Six hundred pg of pre-amplified cDNA were used as input for Tn5 tagmentation, followed by 12 cycles of library amplification with the Nextera XT DNA Library Preparation Kit and IDT for Illumina DNA/RNA UD Indexes, Tagmentation (Illumina, San Diego, USA). After purification with SPRIselect beads (Beckman Coulter, Villepinte, France), library size and concentration were assessed on a Bioanalyzer 2100 (Agilent Technologies, Les Ulis, France). Libraries were sequenced on an Illumina NextSeq 2000 as paired-end 50 bp reads. Image analysis and base calling were performed with RTA 2.7.7 and BCL Convert 3.8.4.

Adapter dimer reads were removed with DimerRemover (−a AGATCGGAAGAGCACACGTCTGAACTCCAGTCAC), and sequencing quality was assessed with FastQC 0.11.5. Reads were aligned to the mouse mm39 genome with STAR v2.7.10b (Dobin et al., 2013), and only uniquely mapped reads were retained. Gene expression was quantified with STAR (−-quantMode GeneCounts). For comparisons between conditions, only genes with more than 50 raw reads were considered.

Differentially expressed genes (DEGs) were identified with DESeq2 (Love et al., 2014) using a p-value < 0.05 (7 dpi) or p-adjusted < 0.05 (uninjured). DEGs were subjected to functional analysis with STRING 12.0 (Szklarczyk et al., 2023) and to pathway analysis with WebGestalt as described above. For heatmaps, expression values were centered and normalized with Cluster 3.0 (De Hoon et al., 2004) and visualized with Morpheus (https://software.broadinstitute.org/morpheus/). Genes were clustered by K-means with gene tree construction, Pearson correlation and average linkage as clustering parameters.

### RNA-seq on whole mouse TA muscles

Total RNA was extracted from injured TA muscles of young and aged mice treated with vehicle or prednisone at 7 dpi, using TRIzol reagent (TRI Reagent, Molecular Research Center, TR-118), and RNA integrity was assessed on a Bioanalyzer. Libraries were prepared and sequenced by the GenomEast platform (IGBMC) according to the standard Illumina protocol (HiSeq 4000, single-end 50 bp reads). Image analysis and base calling were performed with RTA 2.7.7 and bcl2fastq 2.17.1.14. Adapter dimer reads were removed with DimerRemover, and sequencing quality was assessed with FastQC0.11.2. Reads were mapped to the mouse mm10 genome with STAR v2.7.10b (Dobin et al., 2013), and only uniquely aligned reads were retained. Gene expression was quantified with HTSeq 0.11.0 (Anders et al., 2015), and only genes with more than 100 raw reads were considered. DEGs were identified with DESeq2 (Love et al., 2014) using a p-value < 0.05 and analyzed as described for SMART-seq.

### Cell culture and human myoblast analyses

LHCN-M2 immortalized human myoblasts (Zhu et al., 2007), provided by J. Laporte, IGBMC) were grown in a 4:1 mixture of Dulbecco’s modified Eagle medium (DMEM, 4.5 g/l glucose; Invitrogen, 41965-062) and Medium 199 (Invitrogen, 22340-020) supplemented with 20% fetal calf serum (FCS), fetuin (25 µg/ml), human insulin (5 µg/ml), human EGF (5 ng/ml), human bFGF (0.5 ng/ml), gentamicin (40 µg/ml) and dexamethasone (0.2 µg/ml) (medium prepared by the PluriCell East platform, IGBMC). Cells were maintained at 37 °C in a humidified atmosphere with 5% CO_2_.

### siRNA transfection

Because the growth medium contains dexamethasone, cells were switched to charcoal-treated medium 24 h before transfection. At 70% confluence, cells were transfected with 25 nM siRNA against GR (5?-GCUUUGCUCCUGAUCUGAUUAUUAA-3?) or a scrambled control (5?-AGGUUCCGUGUACGUAAGACAAACU-3?) (Invitrogen) using Lipofectamine RNAiMAX (Invitrogen) according to the manufacturer’s instructions. Transfection was performed in charcoal-treated growth medium, and cells were harvested 3 days later.

### Immunocytofluorescence

Cells dedicated to immunofluorescence were seeded on coverslips. At 70-80% confluence, cells were washed twice with 1X PBS, fixed for 15 min in 4% PFA, washed and permeabilized for 10 min in PBS containing 0.1% Triton X-100. After three washes in 1X PBS, cells were blocked for 1 h at room temperature in PBS-Tween containing goat serum (PBS-T SG). Cells were washed again three times and incubated overnight at 4 °C in PBS-T SG with primary antibodies against GR (Proteintech, 24050-1-AP), β-actin (Santa Cruz Biotechnology, sc-47778) and KI67 (Invitrogen, 14-5698-82), all at 1:500. After three washes, cells were incubated for 2 h at room temperature with secondary antibodies (1:500) and mounted in DAPI-Fluoromount-G (Invitrogen, 00-4959-52). KI67^+^ nuclei were counted manually.

### RNA extraction from cultured human myoblasts

LHCN-M2 cells were homogenized in NucleoZOL reagent (500 µl per well of 6-well plates) (MACHEREY-NAGEL, Germany). RNA was isolated using a standard isopropanol extraction protocol, and quantified by spectrophotometry (Nanodrop, Thermo Fisher). RNA was diluted in 20 µl of RNase-free water and stored at −80 °C.

### Reverse transcription and quantitative PCR (qPCR)

Total RNA (2 µg) was reverse transcribed with SuperScript IV (Invitrogen, 18090010) and oligo(dT) primers. RNA was diluted in nuclease-free water to a final volume of 11 µl, and 1 µl of 10 mM dNTPs (Roche Diagnostics, 04728858001) and 1 µl of 50 µM oligo(dT) primers (Invitrogen) were added. Samples were heated for 5 min at 65 °C and cooled to 4 °C for 5 min. A second mix containing 4 µl of 5X SSIV buffer, 1 µl of 100 mM dithiothreitol (DTT), 1 µl of H_2_O and 1 µl of SuperScript IV reverse transcriptase (all from Invitrogen, 18090010) was then added. Samples were incubated for 10 min at 23 °C, 10 min at 55 °C and 10 min at 80 °C, then held at 4 °C. cDNA was diluted 1:10 in nuclease-free water.

Quantitative PCR (qPCR) was performed with the SYBR® Green PCR kit (Roche) according to the manufacturer’s protocol. Each 10 µl reaction contained 2 µl cDNA, 5 µl 2X SYBR Green master mix, 1 µl primer mix (3 µM each) and 2 µl H_2_O. Reactions were run on a CFX Opus 384 Real-Time PCR System (Bio-Rad) with a pre-incubation of 5 min at 95 °C followed by 45 cycles of 15 s at 95 °C, 20 s at 62 °C and 15 s at 72 °C. Primers are listed in Table S3. *RPLP0* was used as the reference gene, and data were analyzed with the ΔΔCt method. Primer efficiency (E) was calculated from the slope of a standard curve as E = (10^(−1/slope)^ - 1) X 100.

### Protein analysis

TA muscles were mechanically ground in RIPA buffer [50 mM Tris (pH 7.5), 1% NP-40, 0.5% sodium deoxycholate, 0.1X SDS, 150 mM NaCl, 5 mM EDTA and a protease inhibitor cocktail (cOmplete, Roche, 11873580001) used at 1X at 4 °C. Alternatively, adherent cells were harvested in RIPA buffer and kept on ice for 10 min. TA muscles and cell lysates were centrifuged at 12,000 g for 10 min at 4°C, and the supernatants were collected for further analysis.

Protein concentration was determined with a Bradford assay, and 20-30 µg of protein per lane were separated on 8% polyacrylamide gels and transferred to Hybond nitrocellulose membranes (Amersham Biosciences). Membranes were blocked for 1 h at room temperature in PBS-Tween containing 5% non-fat dry milk, then probed with specific antibodies targeting GR (Proteintech, 24050-1-AP, 1:500), α-TUBULIN (IGBMC, 1Tub2A2, 1:5000), GAPDH (Cell Signaling, #2118, 1:5000) and actine (homemade, 1:5000).

Secondary antibodies conjugated to horseradish peroxidase (anti-mouse-HRP, Cell Signaling, 7076S, 1:5000; anti-rabbit-HRP, Cell Signaling, 7074S, 1:5000) were detected using an ImageQuant 800 chemiluminescence detection system (GE Healthcare, Amersham). Proteins were quantified with Fiji (ImageJ).

### Statistical analyses

Data processing and analysis were conducted using DESeq2, Microsoft Excel and GraphPad Prism 9. Data are presented as mean ± SEM. Unless otherwise stated, n refers to the number of individual mice (in vivo experiments) or to the number of independent biological replicates (cell culture experiments). Two groups were compared with an unpaired two-tailed Student’s t-test, or with a Mann-Whitney test when normality was rejected; more than two groups were compared with a one-way or two-way ANOVA followed by Tukey’s post hoc test. The specific test applied is indicated in each figure legend. Statistical significance was defined as follows: ns, p > 0.05 (non-significant); *, p < 0.05; **, p < 0.01; ***, p < 0.001; and ****, p < 0.0001.

## Acknowledgments

We thank Dr. J. Laporte (IGBMC) for generously providing the LHCN-M2 cell line. We are grateful to Brayann Weiss for his valuable technical support, and to Zakary Haili and Ludivine Dulac for helpful assistance. We extend our gratitude to Muriel Philipps from the flow cytometry IGBMC facility. We are grateful to Prof. G. Kardon for the *Pax7*-CreER^T2^ line. We acknowledge the support of the IGBMC core facilities, including the animal facility, photonic and electron microscopy platforms, cell culture service, flow cytometry, and histopathology services, as well as the Mouse Clinical Institute (ICS, Illkirch-Graffenstaden, France) and GenomEast facility, a member of the ‘France Génomique’ consortium (ANR-10-INBS-0009). We are grateful to Jan Tuckerman, Bénédicte Chazaud and Rémi Mounier for helpful discussions.

## Author contributions

D.D., S.S.C., D.M., and E.C. formulated the initial hypothesis. E.C., S.S.C., J.G.R., Q.C., A.Bi., and A.Bo. carried out functional, molecular, and histological experiments. E.C. and R.L. performed mouse experiments. D.D., T.Y., C.K. and E.C. executed bioinformatics analyses. E.C. and E.G. took responsibility for imaging analyses. E.C., S.S.C. and D.D. took primary responsibility for data analysis and writing the manuscript.

## Ethical approval

The animal study protocols have been approved by the French Research Ministry (MESR) according to the 2010/63/EU directive under the numbers 48090-2024022308468680 and 23477-2019121617036403.

## Declaration of generative AI and AI-assisted technologies

None declared.

## Data availability

Custom macro for CSA measurements quantification is available on GitLab (CSA: the BIOP cellpose extension guide).

