## Supplementary figures and tables for "Satellite cell glucocorticoid receptor restrains myogenic commitment without mediating the effects of therapeutic glucocorticoids on muscle repair"

This file includes Supplementary Figures 1-6 and Supplementary Tables 1-3

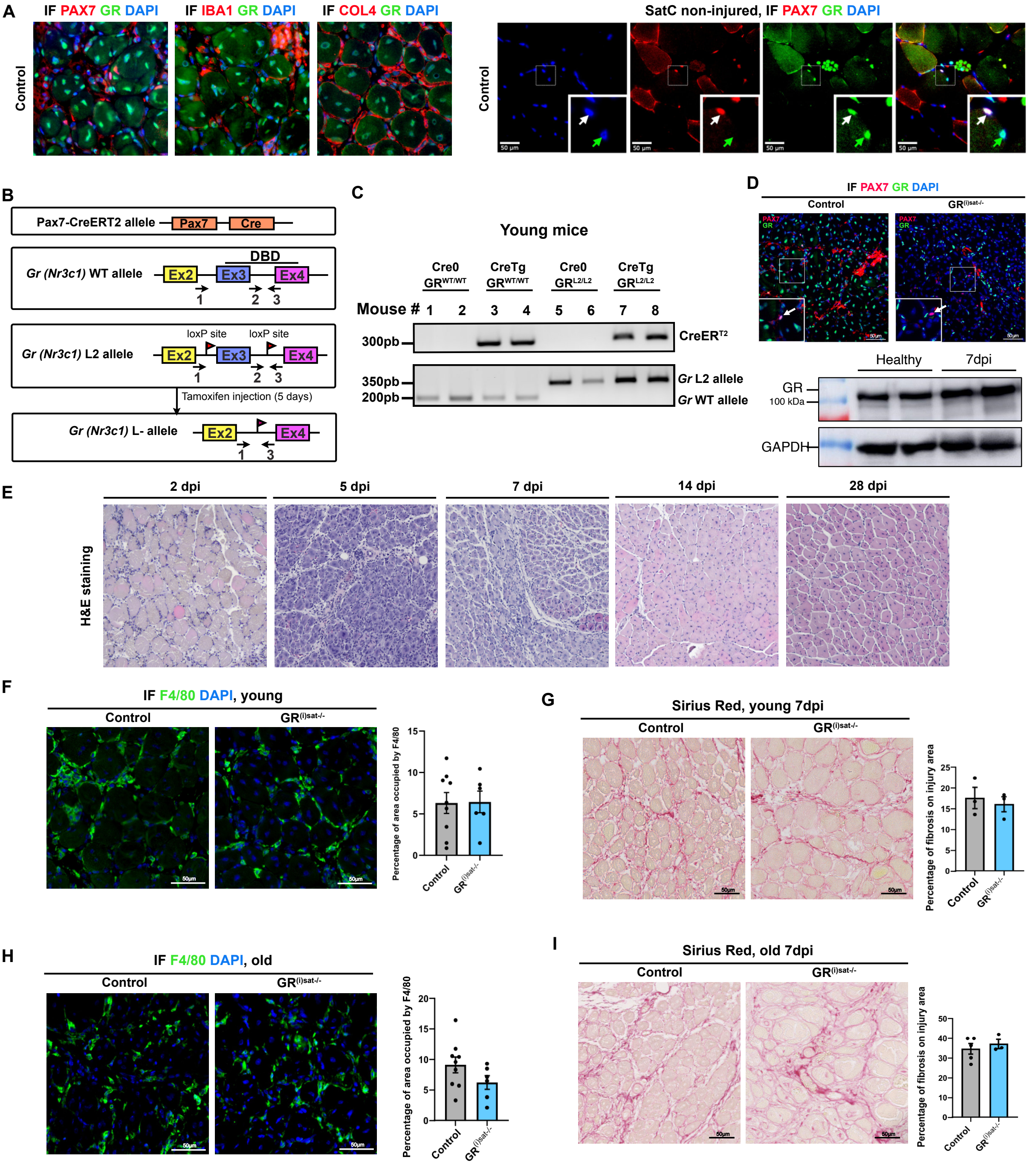


**Figure S1: GR expression in muscle and generation of GR$^{(i)sat-/-}$ mice**

**(A)** Immunofluorescent detection of GR (green) and PAX7, IBA1 or COL4 (red) on TA sections from control mice at 7 dpi. Right, PAX7 (red) and GR (green) in SatC from uninjured muscle. Nuclei counterstained with DAPI. Scale bars, 25 and 50 µm.

**(B)** Schematic representation of the *Pax7*-CreER$^{T2}$, wild-type (WT), floxed (L2) and CreER$^{T2}$-mediated exon 1 deleted (L-) GR alleles. LoxP sites are shown by arrowheads. Primers used for allele characterization are depicted by arrows and sequences are in Table S1

**(C)** Representative electrophoresis of PCR-amplified genomic DNA from tail biopsy of young mice, illustrating the CreER$^{T2}$ transgene (top) and the *Nr3c1* L2 and wild-type (WT) alleles (bottom), with distinct amplicon sizes: CreER$^{T2}$ (300 bp), L2 (360 bp) and WT (200 bp).

**(D)** Immunofluorescent detection of PAX7 (red) and GR (green) on TA sections from control and GR$^{(i)sat-/-}$ mice, with insets showing individual SatC. Scale bars, 50 µm. Below, western blot of GR in TA muscle from uninjured and 7 dpi control mice, with GAPDH as loading control.

**(E)** Representative H&E staining of TA sections from control mice at 2, 5, 7, 14 and 28 dpi after cardiotoxin injury.

**(F)** Immunofluorescent detection of F4/80 and corresponding quantification of the area occupied by macrophages in young control and GR$^{(i)sat-/-}$ mice at 7 dpi. Scale bars, 50 µm. Data are mean ± SEM. Statistical test used was two-tailed Mann-Whitney test.

**(G)** Sirius Red staining and corresponding quantification of fibrosis over the injured area in young control and GR$^{(i)sat-/-}$ mice at 7 dpi. Scale bars, 50 µm. Data are mean ± SEM. Statistical test used was two-tailed Mann-Whitney test.

**(H, I)** Immunofluorescent detection of F4/80 (H) and Sirius Red staining (I), with corresponding quantification, in aged control and GR$^{(i)sat-/-}$ mice at 7 dpi. Scale bars, 50 µm. Data are mean ± SEM. Statistical test used was two-tailed Mann-Whitney test.

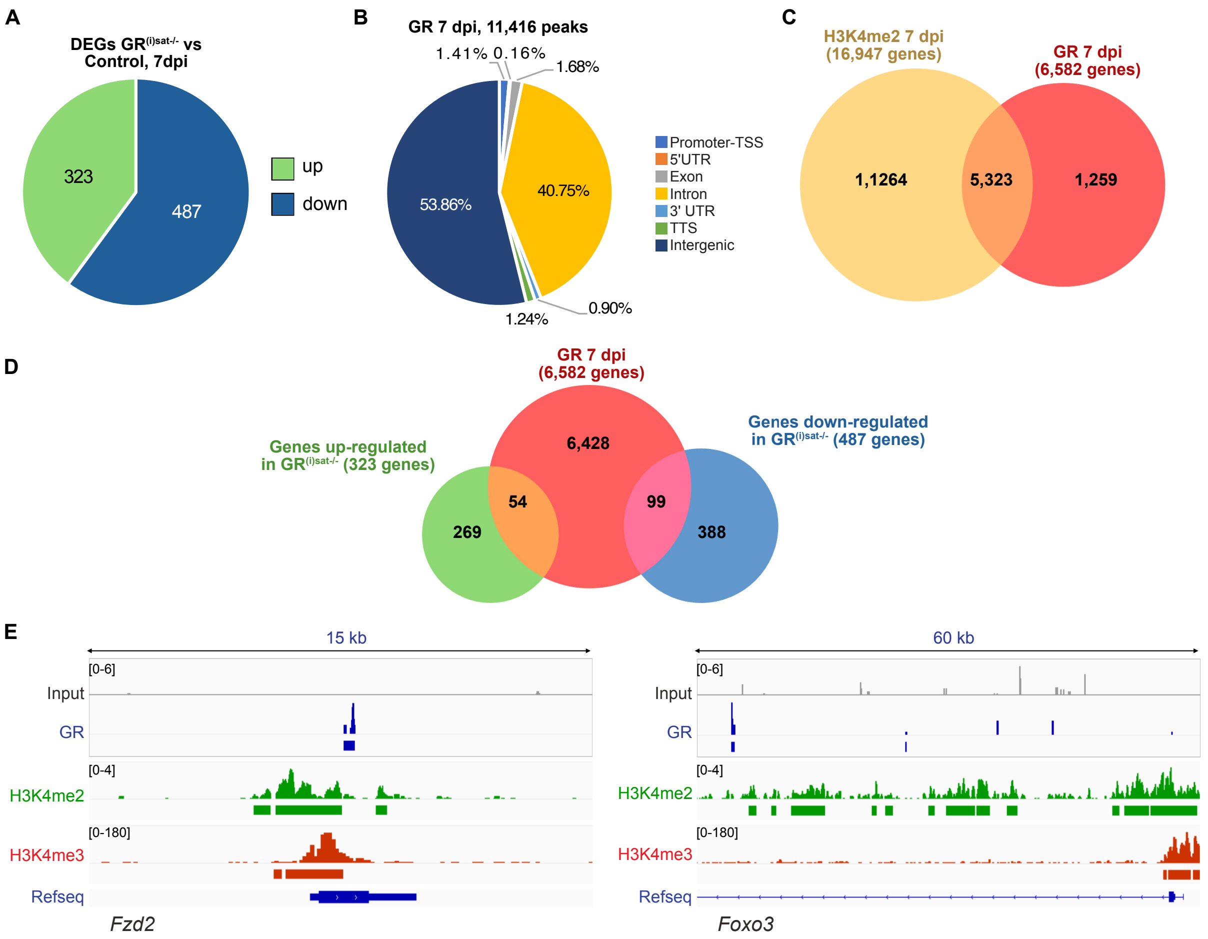


**Figure S2: GR binding in satellite cells at 7 dpi**

**(A)** Pie chart depicting the number of down- (487) and up-regulated (323) genes in GR$^{(i)sat-/-}$ versus control SatC FACS-isolated from 7 dpi TA of young male mice.

**(B)** Pie chart depicting the genomic location of the 11,416 GR binding sites identified by CUT&RUN in SatC FACS-isolated from 7 dpi TA of young male mice.

**(C)** Overlap between genes bound by GR (6,582 genes) and genes marked by H3K4me2 peaks in SatC from 7 dpi TA.

**(D)** Overlap between genes bound by GR (6,582 genes) and genes down- (487) or up-regulated (323) in GR$^{(i)sat-/-}$ SatC from 7 dpi TA.

**(E)** GR, H3K4me2 and H3K4me3 localization at the *Fzd2* and *Foxo3* loci in SatC from 7 dpi TA.

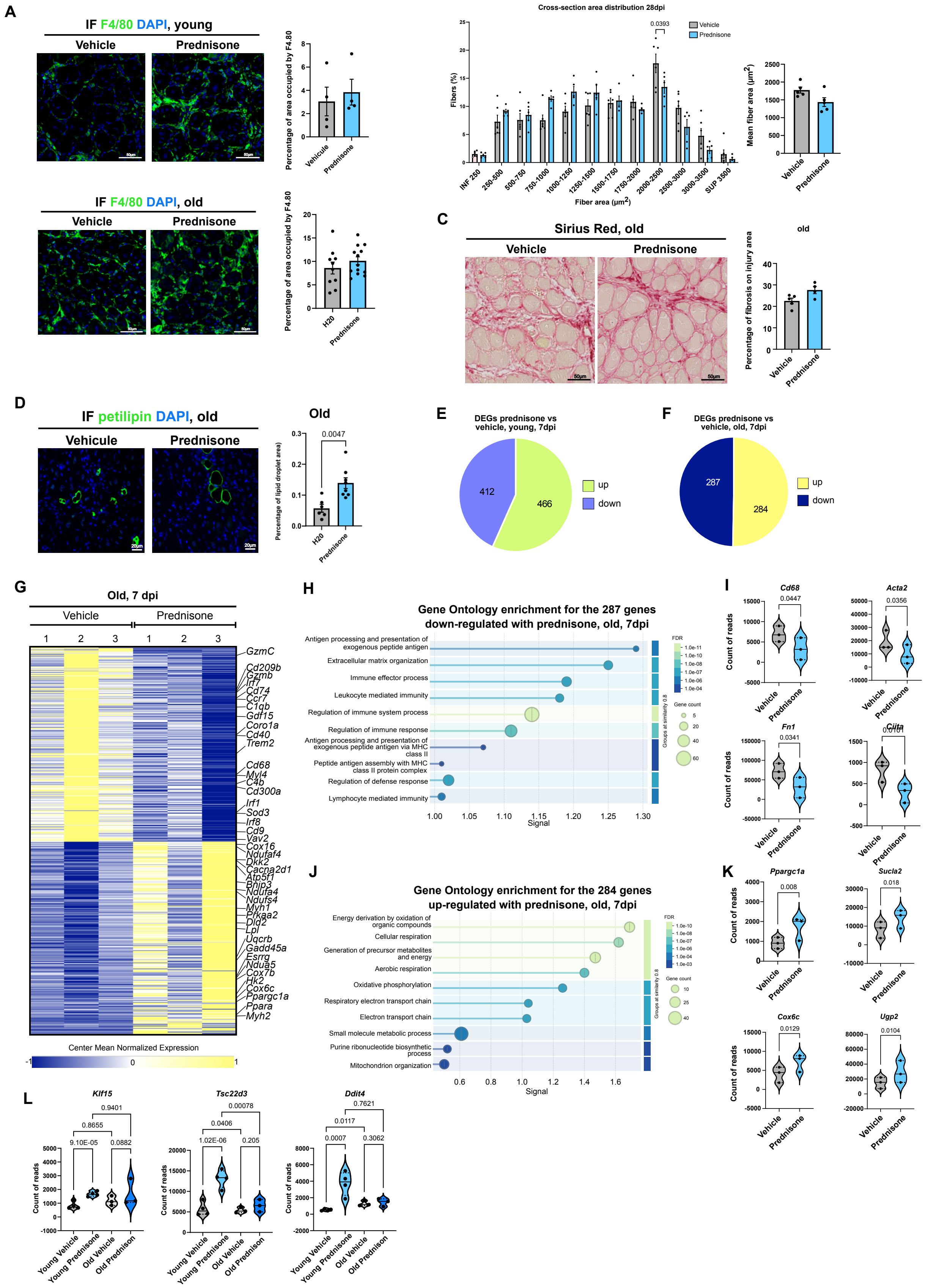


**Figure S3: Effects of chronic prednisone in young and aged mice**

**(A)** Representative immunofluorescent labeling of F4/80 and corresponding quantification of the area occupied by macrophages, in 7 dpi TA of young (top) and aged (bottom) mice treated with vehicle or prednisone. Nuclei were stained with DAPI. Scale bars, 50 µm. Data are presented as mean ± SEM. Statistical test used was two-tailed Mann-Whitney test.

**(B)** Distribution of myofiber cross-sectional area (CSA) and mean fiber area in 28 dpi TA of young mice treated with vehicle or prednisone. Data are presented as mean ± SEM. Statistical test used was two-tailed Mann-Whitney test.

**(C)** Representative Sirius Red staining and corresponding quantification of fibrosis over the injured area, in 7 dpi TA of aged mice treated with vehicle or prednisone. Scale bars, 50 µm. Data are presented as mean ± SEM. Statistical test used was two-tailed Mann-Whitney test.

**(D)** Representative immunofluorescent labeling of perilipin and corresponding quantification of lipid droplet area in 7 dpi TA of aged mice treated with vehicle or prednisone. Nuclei were stained with DAPI. Scale bars, 20 µm. Data are presented as mean ± SEM. Statistical test used was two-tailed Mann-Whitney test.

**(E-F)** Pie charts depicting the number of down- and up-regulated genes in 7 dpi TA of young (A: 412 down, 466 up) and aged (B: 287 down, 284 up) mice treated with prednisone versus vehicle.

**(G)** Heatmap showing the mean-centered normalized expression of differentially expressed genes in 7 dpi TA of aged mice treated with prednisone versus vehicle.

**(H-K)** Gene ontology enrichment of the 287 down- (H) and 284 up-regulated (J) genes, and normalized read counts of representative down- (I) and up-regulated (K) genes, in 7 dpi TA of aged mice. P-values were obtained from DESeq2.

**(L)** Normalized read counts of *Klf15*, *Tsc22d3* and *Ddit4* in 7 dpi TA of young and aged mice treated with vehicle or prednisone. P-values were obtained from DESeq2.

Prednisone was administered daily at 1 mg/kg in young and 0.1 mg/kg in aged mice, from 7 days before injury until 7 dpi.

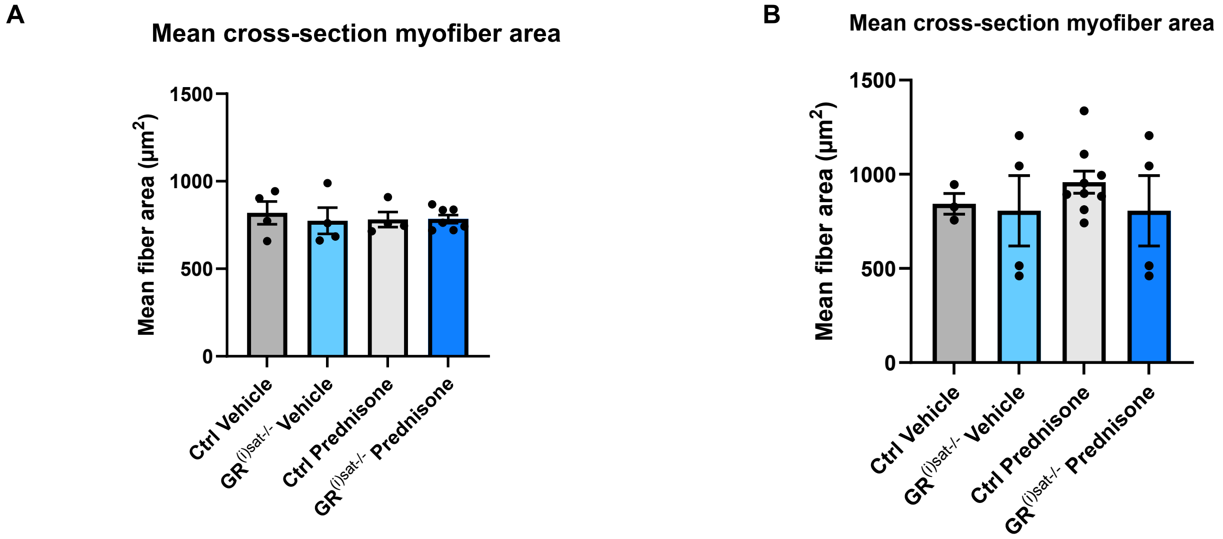


**Figure S4: Mean myofiber area in control and $GR^{(i)sat-/-}$ mice under chronic prednisone**

(**A, B**) Mean myofiber cross-sectional area at 7 dpi in young (**A**) and aged (**B**) control and $GR^{(i)sat-/-}$ mice treated with vehicle or prednisone. Data are mean ± SEM. Statistical test used is Kruskal-Wallis.

Prednisone was administered daily at 1 mg/kg in young and 0.1 mg/kg in aged mice, from 7 days before injury until 7 dpi.

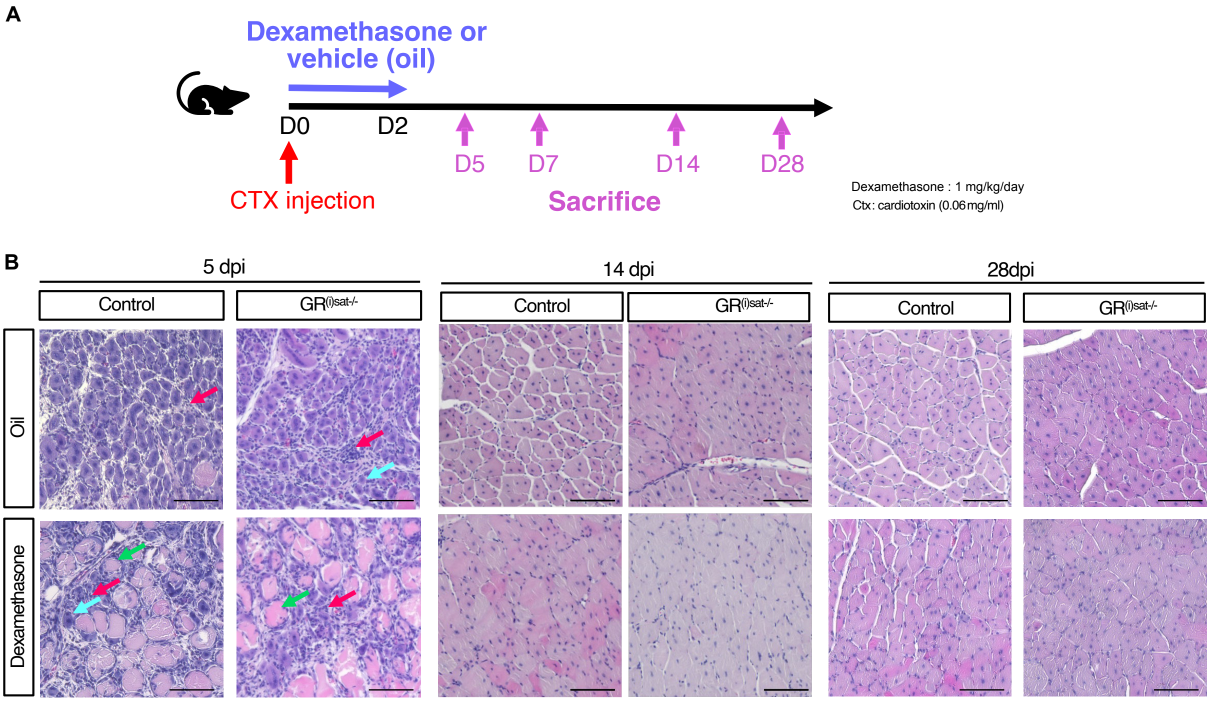


**Figure S5: Early dexamethasone treatment and later stages of regeneration**

**(A)** Schematic representation of the experimental design. Young control and GR$^{(i)sat-/-}$ mice received intraperitoneal injections of dexamethasone (1 mg/kg/day) or vehicle (sunflower oil containing 1.5% EtOH) immediately after cardiotoxin injury (0.06 mg/ml) of the TA (day 0), then at 1 and 2 dpi. TA muscles were collected at 5, 7, 14 and 28 dpi.

**(B)** Representative H&E staining of TA muscles of control and GR$^{(i)sat-/-}$ mice treated with vehicle or dexamethasone, at 5, 14 and 28 dpi. Cyan arrows indicate newly formed fibers, green arrows necrotic fibers and red arrows inflammatory infiltrate. Scale bars, 50 µm.

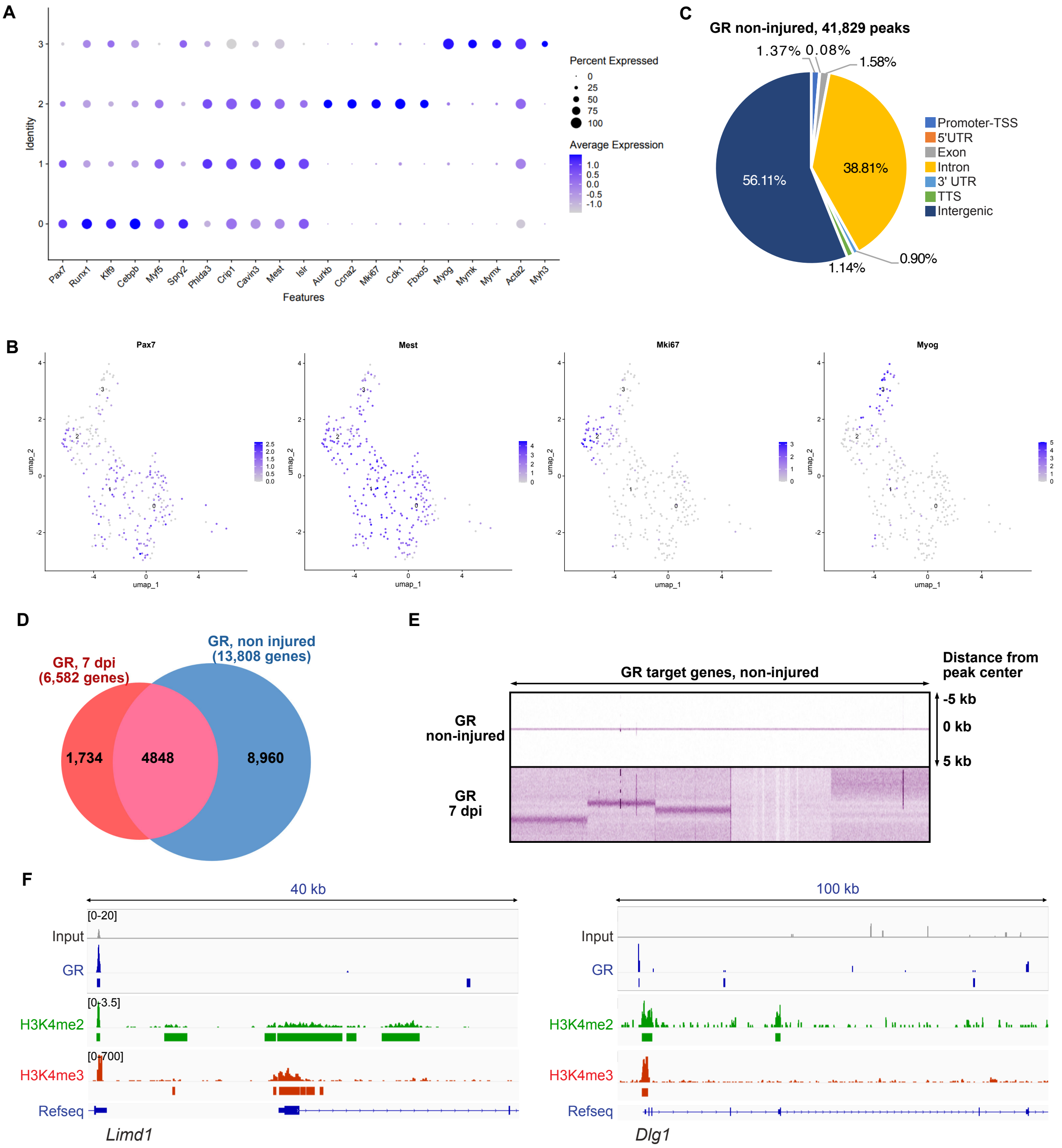


**Figure S6: Satellite cell states and GR occupancy in uninjured muscle**

**(A)** Dot plot heatmap of selected quiescence, proliferation and differentiation marker genes in satellite cell reclusters (clusters 0-3; 0 = quiescent, 1 = activated, 2 = dividing, 3 = differentiated), selected from scRNA-seq analysis performed in 5 dpi TA (Rizk et al., 2026). Dot size represents the percentage of cells in each cluster in which the gene is detected, and color represents the average scaled expression, from light grey (low) to blue (high).

**(B)** Feature plots representing *Pax7*, *Mest*, *Mki67* and *Myog* expression levels on the UMAP of satellite cell reclusters (clusters 0-3) from 5 dpi TA. Each dot represents one cell, colored according to its normalized expression level, from light grey (low) to blue (high).

**(C)** Pie chart depicting the genomic location of the 41,829 GR binding sites identified by CUT&RUN in SatC FACS-isolated from uninjured muscles of young mice.

**(D)** Overlap between genes bound by GR in SatC from uninjured muscles (13,808 genes) and from 7 dpi TA (6,582 genes).

**(E)** Heatmap of GR CUT&RUN signal in SatC from uninjured muscles and from 7 dpi TA, centered on GR peaks identified in uninjured SatC.

**(F)** GR, H3K4me2 and H3K4me3 localization at the *Limd1* and *Dlg1* loci in SatC from uninjured muscles.

**Table S1.** Primers used for mouse genotyping.

| Allele | Forward primer (5′-3′) | Reverse primer (5′-3′) | Product size |
|---|---|---|---|
| *Pax7*-CreER$^{T2}$ | TTCCCGCAGAACCTGAAGATGTTCG | GGGTGTTATAAGCAATCCCCAGAAATGC | 300 bp |
| *Nr3c1* L2 / WT | GACTGACAAAATCAGTGACCCTGGG | GTCAACACATGATCACCTTGCAGTC | 350 bp (L2) 200 bp (WT) |

**Table S2.** Antibodies and reagents for immunofluorescence on muscle sections

| Target / reagent | Conjugate | Supplier | Catalogue no. | Dilution |
|---|---|---|---|---|
| PAX7 | | Thermo Fisher | PA1-117 | 1/1000 |
| GR | | Proteintech | 24050-1-AP | 1/500 |
| F4/80 | | Cell Signaling | 70076S | 1:1000 |
| Perilipin | | Abcam | ab3526 | 1:200 |
| Rabbit IgG | | PeproTech | 500-P00 | 1:500 |
| Mouse IgG | | Santa Cruz | sc-2025 | 1:500 |
| WGA | Alexa Fluor 555 | Invitrogen | W32464 | 1:500 |
| Goat anti-rabbit IgG | Alexa Fluor 488 | Invitrogen | A11008 | 1:500 |
| Goat anti-rabbit IgG | Alexa Fluor Plus 555 | Invitrogen | A32732 | 1:500 |

**Table S3.** Human primer sequences used for RT-qPCR.

| Gene | Forward primer (5′-3′) | Reverse primer (5′-3′) |
|---|---|---|
| *NR3C1* (GR) | ATAGCTCTGTTCCAGACTCAACT | TCCTGAAACCTGGTATTGCCT |
| *DDIT4* | CTTCTACGCTCCGGCACTCT | CCAGGTAAGCCGTGTCTTCCTCC |
| *TRIM63* (MURF1) | GTCGAGTGACCAAGGAGAACA | CAGGGACTGGATGGCAGTTT |
| *MKI67* | TACTATCAAAAGGAGCGGGGTC | TCTTGAGCTTTCTCATCAGGG |
| *RUNX1B* | TCTGCAGAACTTTCCAGTCG | GTCGGGGAGTAGGTGAAGG |
| *RUNX1* | CCACCTACCACAGAGCCATCAA | TTCACTGAGCCGCTCGGAAAAG |
| *FOXO1* | GAGGGTTAGTGAGCAGGTTACA | CTGCACACATTGGGCAAACA |
| *MYH3* (eMHC) | AACTGGAGACCAGGATCCGA | TCCTTGACCCTCCGCTCATA |
| *MYOG* | CGCCAACCCAGGGGATCA | AGGCCCCTGCTACAGAAGTA |
| *RPLP0* | CGTCCTCGTGGAAGTGACAT | TAGTTGGACTTCCAGGTCGC |
